# Mammalian Orthoreovirus Factories Modulate Stress Granule Protein Localization by Interaction with G3BP1

**DOI:** 10.1101/169433

**Authors:** Promisree Choudhury, Luke Bussiere, Cathy L. Miller

## Abstract

Mammalian orthoreovirus (MRV) infection induces phosphorylation of translation initiation factor eIF2α which promotes formation of discrete cytoplasmic inclusions, termed stress granules (SGs). SGs are emerging as a component of the innate immune response to virus infection, and modulation of SG assembly is a common mechanism employed by viruses to counter this antiviral response. We previously showed that MRV infection induces SGs early, then interferes with SG formation as infection proceeds. In this work, we found that SG associated proteins localized to the periphery of virus-encoded cytoplasmic structures, termed virus factories (VFs), where viral transcription, translation, and replication occur. The localization of SG proteins to VFs was dependent on polysome dissociation and occurred via association of SG effector protein, G3BP1, with MRV non-structural protein σNS, which localizes to VFs via association with VF nucleating protein, μNS. Deletion analysis of the σNS RNA binding domain and G3BP1 RNA (RRM) and ribosomal (RGG) binding domains showed that the association and VF localization of G3BP1 is not occurring solely through RNA or ribosomal binding, but requires both RNA and ribosomal binding domains of G3BP1 for maximal VFL localization and σNS association. Co-expression of σNS and μNS resulted in disruption of normal SG puncta, and in cells lacking G3BP1, MRV replication was enhanced in a manner correlating with strain-dependent induction of host translation shutoff. These results suggest that σNS association with and relocalization of G3BP1 to the VF periphery plays a role in SG disruption to facilitate MRV replication in the host translational shutoff environment.

**IMPORTANCE:** SGs and SG effector proteins have emerged as important, yet poorly understood, players in the host’s innate immune response to virus infection. MRV infection induces SGs early during infection that are dispersed and/or prevented from forming during late stages of infection despite continued activation of the eIF2α signaling pathway. Cellular and viral components involved in disruption of SGs during late stages of MRV infection remain to be elucidated. This work provides evidence that MRV disruption of SGs may be facilitated by association of MRV non-structural protein σNS with major SG effector protein G3BP1 and subsequent localization of G3BP1 and other SG associated proteins around the periphery of virus encoded factories, interrupting the normal formation of SGs. Our findings also reveal the importance of G3BP1 as an inhibitor of MRV replication during infection for the first time.

## INTRODUCTION

MRV infection triggers the host innate immune response in part by activation of protein kinase R (PKR) resulting in phosphorylation of the alpha subunit of eukaryotic translation initiation factor 2 (eIF2α) (1). eIF2α phosphorylation leads to host translational shutoff inducing the formation of cytoplasmic RNA-protein granules called stress granules (SGs) (2). SGs are composed of translationally silent mRNAs, translation initiation factors, stalled ribosomal subunits, and RNA regulatory proteins (3). To date, the mechanism of SG formation is not fully understood and over 100 genes have been suggested to be involved in the process of SG assembly and disassembly (4). Studies have identified several proteins as critical for nucleating SGs, including T-cell intracellular antigen 1 (TIA-1), TIA-1-related protein (TIAR), and Ras-GAP SH3-binding protein1 (G3BP1) (4). G3BP1 is a ubiquitously expressed cytosolic protein that was originally identified as a binding partner of Ras GTPase activating protein (GAP) (5), although that binding has been recently brought into question (6). Overexpression of G3BP1 induces SG formation, whereas expression of a central domain of the protein inhibits SG formation (7). This central domain contains a serine at residue 149, that when dephosphorylated, induces SG formation (7). G3BP1 also interacts with the ubiquitin specific protease USP10 (8), and the cytoplasmic activation- and proliferation-associated protein 1 (Caprin1) (9), and this binding appears to modulate SG formation and PKR activation (10, 11). G3BP2, a homolog of G3BP1 is also recruited to SGs (12).

Recently, studies have suggested that SGs, or structurally similar antiviral stress granules (avSGs), serve as a downstream component of the innate immune response to virus infection, acting as a platform for recognition of non-self (viral) RNA and subsequent activation of important immune modulators including PKR and retinoic acid inducible gene I, (RIG-I) (13-15). Many viruses have been shown to modulate SG composition, assembly, or disassembly to facilitate their replication and infection cycle (16, 17). For example, poliovirus induces SG formation at early times in infection and disrupts SGs during later stages of infection by cleavage of G3BP1 by the poliovirus 3C proteinase (18). Infection with Semliki Forest virus or rotavirus results in host translational shutoff via phosphorylation of eIF2α, and subsequent SG formation at early phases of infection with disruption of the granules by unknown mechanisms during the later stages of infection (19, 20). In contrast, the respiratory syncytial virus induction of SGs appears to facilitate viral replication (21).

Mammalian orthoreovirus (MRV) is the type species of the orthoreovirus genus of the double-stranded RNA (dsRNA) family *Reoviridae.* MRV is a clinically benign virus in immune-competent hosts, making it an ideal tool to model various aspects of virus infection. Moreover, MRV has inherent oncolytic properties, and is thus being investigated as a novel cancer therapy in both pre-clinical and clinical settings (22, 23). Early during MRV infection, viral factories (VFs) are formed which originate as small punctate inclusions spread throughout the cytoplasm, which grow in size and migrate to the perinuclear region of the cell as infection proceeds (24, 25). VFs are suggested to be the primary sites for virus transcription, replication, and assembly (26-28) and recent studies have indicated that they also function as the primary location for viral protein synthesis by interacting with cellular translation initiation factors and components of the 43S pre-initiation ribosomal complexes at the factory margins (29). The MRV non-structural protein μNS forms the structural matrix of VFs, and when expressed in the absence of other proteins, forms structures very similar to VFs, termed viral factory-like structures (VFLs) (25). Moreover, when expressed with μNS in cells, viral proteins σNS, σ2, μ2, λ1, λ2, and λ3 are recruited to VFLs, suggesting that μNS is also involved in the recruitment or retention of these proteins to VFs in infected cells (25, 30-33).

We have previously shown that SGs are formed at early times following MRV infection in an eIF2α phosphorylation-dependent manner. MRV-induced SG formation is dependent on virus uncoating but does not require virus transcription or translation (34). However, as MRV infection proceeds, SGs are disrupted, in a manner requiring MRV transcription and/or translation, suggesting that the virus encodes a gene product involved in dissolution of SGs. Moreover, at late times in MRV infection in most infected cells, SG formation is prevented even in the presence of phosphorylated eIF2α induced by the virus or other stress inducers, such as sodium arsenite, suggesting that MRV disrupts a signal between eIF2α phosphorylation and SG formation. In addition, the presence of SGs in cells results in the inhibition of virus and host translation whereas MRV-induced disruption of SGs correlates with the release of viral, but not cellular, mRNAs from translational inhibition even in the presence of phosphorylated eIF2α (35). Taken together, these studies suggest that the host responds to MRV infection by forming SGs to interfere with translation, however, the virus can overcome this translational inhibition in a manner that correlates with virus-dependent SG disruption. Although the formation and disruption of SGs during MRV infection are well-established, the role played by the virus in regulating the process remains to be elucidated. An interaction between the VF matrix forming protein μNS and SGs induced by sodium arsenite was previously identified, suggesting MRV proteins may interact with SG proteins to modulate SGs (36). In this study, we set out to identify specific SG effector proteins that may be modulated by MRV to disrupt SGs, and to identify viral proteins involved in this process. We found that MRV infection alters the localization of SG-associated proteins to the periphery of VFs in a subset of infected cells, and that this occurs as a result of the virus non-structural protein, σNS, binding to both G3BP1 and the VFL-forming viral protein μNS. The C-terminus of G3BP1, which contains an RNA recognition motif and arginine/glycine rich motif (RRM and RGG) are involved in this interaction and G3BP1 mutants lacking these domains exhibit significantly reduced localization to VFLs. We additionally provide evidence that co-expression of MRV proteins σNS and μNS prevents canonical SG formation and G3BP1 expression is detrimental to MRV replication in a strain specific manner that strongly correlates with virus induced host translational shutoff.

## MATERIALS AND METHODS

### Cells, viruses, and antibodies

HeLa (human cervical cancer cells), Cos-7 (monkey kidney fibroblast), wildtype or G3BP1 -/- MEF (mouse embryonic fibroblast) (11, 37) and wildtype or ΔG3BP1 or ΔΔG3BP1/2 U2OS (human bone osteosarcoma) (10) cells were maintained in Dulbecco’s modified Eagle’s medium (DMEM) (Life Technologies) containing 10% fetal bovine serum (Atlanta Biologicals) and penicillin (100 IU/ml)-streptomycin (100 μg/ml) (Mediatech). L929 cells were maintained in Minimum Essential Medium with Joklik Modification (Sigma) containing 2% fetal bovine serum (HyClone Laboratories), 2% bovine calf serum (HyClone Laboratories), penicillin (100 IU/ml)-streptomycin (100 μg/ml) (Mediatech), and L-Glutamine (2 mM; Mediatech). The primary antibodies used are as follows: rabbit polyclonal anti-G3BP1 (α-G3BP1) antibody (Novus Biologicals, NBP1-18922), rabbit polyclonal anti-Caprin1 (α-Caprin1) antibody (Proteintech, 15112-1-AP), rabbit monoclonal anti-USP10 (α-USP10) antibody (Cell signaling, 8501), goat polyclonal anti-TIA-1 (α-TIA-1) antibody (Santa Cruz Biotechnology, sc-1751), goat polyclonal anti-TIAR (α-TIAR) antibody (Santa Cruz Biotechnology, sc-1749), rabbit polyclonal anti-eIF3b (α-eIF3b) antibody (Bethyl Laboratories, A301-760A), rabbit polyclonal anti-EGFP (α-EGFP) antibody (Living Colors full length GFP antibody, Clontech, 632460), mouse monoclonal anti-σNS (3E10) (α-σNS) antibody and mouse polyclonal anti-μNS (α-μNS) antibody (35, 36, 38). The secondary antibodies used are as follows: Alexa 488- or Alexa 594-conjugated donkey anti-rabbit, anti-mouse or anti-goat immunoglobulin G (IgG) antibodies (Invitrogen). All primary antibodies used were tested by Western blot and found to recognize a protein of the correct molecular weight. MRV stocks (T1L, T2J, and T3D) are our laboratory stocks and were propagated on L929 cells and plaque purified in our laboratory as described previously (39). The T3D virus used in this study was the previously designated Cashdollar strain (40). Virions were stored in dialysis buffer (150 mM NaCl, 10 mM Tris, pH 7.4, 10 mM MgCl_2_) at 4°C.

### Plasmid Construction

All MRV genes examined in this study were cloned into the pCI-neo expression vector (Promega). pCI-σNS (T1L, T3D) (32), pCI-μNS (T1L, T3D) (25), pCI-σNS (Δ2-11) (32), p-EGFP/μNS(471-721) (41), and p-EGFP/NSP5 (42) were previously described. p-G3BP1/EGFP was made by amplifying G3BP1 from pCMV6-G3BP1 (Origene, NM_005754) using G3BP1 sequence specific primers flanked on each end with restriction sites for HindIII and EcoR1. The PCR fragment was then digested with HindIII and EcoR1, and inserted into HindIII and EcoR1 digested p-EGFP-C1. To make pCI-G3BP1, the G3BP1 gene was amplified by PCR from p-G3BP1/EGFP using primers with restriction sites for NheI and XmaI at the 5′ and 3′ terminal ends respectively. The PCR fragment was digested with NheI and XmaI and ligated into NheI and XmaI digested pCI-neo. To make p-G3BP1/EGFP/μNS(471-721), the G3BP1 gene from pCI-G3BP1 was amplified by PCR using G3BP1 sequence specific primers flanked on each end with NheI and AgeI restriction sites. p-EGFP/μNS(471-721) and the G3BP1 PCR fragment were then digested with NheI and AgeI, and ligated. p-G3BP1/EGFP/NSP5 was made by digesting plasmids p-σNS/EGFP/NSP5 and p-G3BP1/EGFP/μNS(471-721) with restriction enzymes AgeI and ApaI, and ligating the p-EGFP/NSP5 fragment into the p-G3BP1/EGFP/μNS(471-721) vector replacing the EGFP/μNS(471-721) fragment. The G3BP1 mutant plasmids were constructed by designing gBlocks (IDT DNA) of the full-length G3BP1 gene with F33W, S149A, S149E, ΔRRM (Δ340-416), ΔRGG (Δ428-466) mutations flanked on each end with restriction sites for NheI and XmaI. The gBlocks were digested with NheI and XmaI, and ligated into NheI and XmaI digested pCI-neo. pCI-σNS (T2J) was made by ligating an EcoRI and XbaI digested S3 (T2J) gBlock into EcoR1 and XbaI digested pCI-neo. The M3 gene was amplified from T2J infected cells by RT-PCR using M3 sequence specific primers containing terminal EcoR1 and XbaI restriction sites. The PCR fragment was digested with EcoRI and XbaI and ligated into EcoR1 and XbaI digested pCI-neo to construct pCI-μNS (T2J). Plasmids were screened by restriction digestion and inserts were confirmed by sequencing. The sequences of all primers used are available upon request.

### Transfections and Infections

Cells were seeded onto culture plates the day before transfection/infection. For transfection, 1 μg plasmid DNA and 3 μl of Trans-IT LT1 reagent (Mirus Bio) per 1 μg of DNA, were combined with 100 μl of Opti-MEM serum-free media (Thermofisher) and incubated for 20 min at room temperature, then added dropwise to the cell media. Cells were then incubated overnight at 37°C and subjected to downstream assays. For infection, 1-5 cell infectious units (CIU) of MRV strains T1L, T2J or T3D were used per well and incubated overnight at 37°C. CIU for each cell line was determined as previously described (35).

### Immunofluorescence assay

Cells were seeded onto 12-well (3.8 cm^2^) cell culture plates (Corning Inc.) containing 18-mm-diameter coverslips (Fisherbrand) at a density of 7.5X10^4^ cells/well and then incubated overnight at 37°C. Sodium arsenite (SA) (Sigma-Aldrich) was used where indicated at a final concentration of 0.5 mM for 45 mins before fixing. Cycloheximide (Sigma-Aldrich) was used where indicated at a final concentration of 10 μg/ml for 1hour (h) before fixing. Puromycin (Invitrogen) was used where indicated at a final concentration of 1 μg/ml for 1 h before fixing. At 24 h post transfection or infection (h p.t. or h p.i.), cells were fixed at room temperature for 20 min with 4% paraformaldehyde in phosphate buffered saline (PBS) (137 mM NaCl, 3 mM KCl, 8 mM Na_2_HPO_4_, pH 7.5) and then washed two times with PBS. Fixed cells were permeabilized by incubation with 0.2% Triton X-100 in PBS for 5 min and then washed two times with PBS. Primary and secondary antibodies were diluted in 1% bovine serum albumin in PBS. Following permeabilization, cells were incubated for one hour with primary antibody, washed two times with PBS, and then incubated for an additional hour with secondary antibody. Immunostained cells were washed two additional times with PBS and mounted on slides using ProLong gold antifade reagent with DAPI (4, 6-diamidino-2-phenylindole dihydrochloride) (Life Technologies). Samples were examined with a Zeiss Axiovert 200 inverted microscope equipped with fluorescence optics. For high resolution confocal images, an Olympus Fluoview FV1000 laser scanning confocal microscope equipped with spectral deconvolution hardware and four variable-voltage color lasers was used. Images were prepared using ImageJ (43) or Photoshop and Illustrator software (Adobe Systems). For quantification of localization of SG associated proteins G3BP1, Caprin1, USP10, TIA-1, TIAR and eIF3b to VFs with or without drug treatment, or G3BP1 mutants F33W, S149A, S149E, ΔRRM, ΔRGG and ΔRRM/ΔRGG to VFLs, infected or transfected cells with and without the indicated protein around the periphery of VFs or VFLs were counted, and the percentage of cells with localization to VFs or VFLs relative to total infected or transfected cells was determined. At least 100 cells from three biological replicates were counted and the mean and standard deviation of the mean was determined. Statistical significance was determined as needed using a two-tailed student’s t-test and *p* values were calculated with Microsoft Excel. Any statistical difference within groups for which *p* < 0.05 or *p* < 0.01 or *p* < 0.001 were considered statistically significant are indicated with one (^∗^) or two (^∗∗^) or three (^∗∗∗^) asterisks in the figures respectively.

### Proximity Ligation Assay

Proximity ligation assays were performed using DuoLink PLA Technology (Millipore Sigma) plus and minus probes and detection reagents per manufacturer’s instructions. Briefly, transfected cells were subjected to immunofluorescence assay protocol to the post-wash step following incubation with mouse or rabbit primary antibodies. Mouse minus and rabbit plus probe antibodies were diluted in antibody dilution buffer provided by manufacturer, added to coverslips and incubated for 1 h at 37° C in a humidity chamber. Coverslips were washed 2 X 5 min with Wash Buffer A (0.01 M Tris, 0.15 M NaCl and 0.05% Tween 20), and the ligation reaction (1X ligation buffer and ligase provided by manufacturer, diluted 1:40 in polished water) was added to coverslips and incubated for 45 min at 37° C in a humidity chamber. Coverslips were again washed 2 X 2 min with Wash Buffer A, and the amplification reaction (1X Amplification buffer and polymerase provided by manufacturer, diluted 1:80 in polished water) was added to coverslips and incubated for 60 min at 37° C in a humidity chamber. Coverslips were washed 2 X 10 min in Wash Buffer B (.02 M Tris, .01M NaCl), then washed 1 X 1 min in 0.01X Wash Buffer B and mounted on slides using Prolong Gold antifade reagent with DAPI. PLA slides were imaged using an Olympus Fluoview FV1000 laser scanning confocal microscope. Images were acquired using the same relative locational plane and exposure settings within experiments. For quantification, 7-10 cells from experiments were subjected to the 3D Objects Counter analysis tool in ImageJ using the same threshold measurement for each image. The total pixel area from each image was calculated and divided by the number of cells present to arrive at pixel area/cell for each image in each experimental set. Graphs were created using the box plot tutorial of Peltier’s Tech Charts for Excel. Statistical significance was determined using a two-tailed student’s t-test and*p* values were calculated. Any statistical difference within groups for which *p* < 0.05 or *p* < 0.01 or *p* < 0.001 were considered statistically significant are indicated with one (^∗^) or two (^∗∗^) or three (^∗∗∗^) asterisks in the figures respectively.

### Virus growth assay

Wildtype and G3BP1 -/- MEF cells or wildtype and ΔG3BP1 or ΔΔG3BP1/2 U2OS cells were seeded onto 6-well (9.6 cm^2^) plates (Corning Inc). MEF cells were infected with an MOI of 0.1 and U2OS cells were infected with an MOI of 0.01 of T1L, T2J or T3D and the infection was continued for 24, 48 and 72 hours. Cells were harvested and subjected to three freeze/thaw cycles. The virus-infected lysates were serially diluted (10-fold dilutions) in PBS containing 2 mM MgCl_2_ and titers were determined by standard plaque assay on L929 cells (44). Each experiment was performed for three independent biological replicates and repeated for two experimental replicates and the mean of the six experiments was plotted on a line graph with error bars depicting the standard deviation (SD). Statistical significance was determined using a two-tailed student’s t-test and *p* values were calculated with Microsoft Excel. Any statistical difference within groups for which *p* < 0.05 was considered statistically significant is indicated with an asterisk (∗) in the figures.

## RESULTS

### Stress granule associated proteins are localized around MRV factories in a subset of infected cells

We and others have shown that during infection with MRV, cells initiate a stress response mediated by phosphorylation of eIF2α, leading to SG formation (34, 45). As infection proceeds, although eIF2α phosphorylation remains detectable, SGs are no longer present in infected cells (35). While canonical SGs were not observed in our prior studies, we observed SG associated proteins surrounding the outer periphery of VFs in a small percentage of infected cells, leading us to hypothesize that VFs may play a role in SG modulation during infection. To more carefully investigate this finding, HeLa cells were infected with MRV strain T2J and at 24 h p.i., cells were fixed and stained with antibodies against the VF-localized MRV non-structural proteins σNS or μNS and G3BP1, a well-characterized SG-associated protein, followed by Alexa fluor-labeled secondary antibodies. The cells were then examined by immunofluorescence microscopy to determine the localization of G3BP1. Similar to what we had previously observed, we found that in most infected cells, G3BP1 displayed a diffuse phenotype, however, in a subset of cells (20-40%, depending on cell type), G3BP1 was found localized around MRV factories (Fig. 1A, first row, Fig. 1C). This phenotype was also observed in Cos-7 (Fig. 1A, second row, Fig. 1C), and MEF cells (Fig. 1A, third row, Fig. 1C), suggesting it is not cell type specific. G3BP1 localization around the outer periphery of VFs was confirmed by high resolution confocal microscopy, and was apparent in all planes of 3D Z-stacks (Fig. 1B and Supplemental movie 1). Similar localization of G3BP1 was observed in T1L and T3D infected HeLa, Cos-7 and MEF cells albeit to a substantially lower level compared to T2J infection (Fig. 1C).

**Fig. 1.**
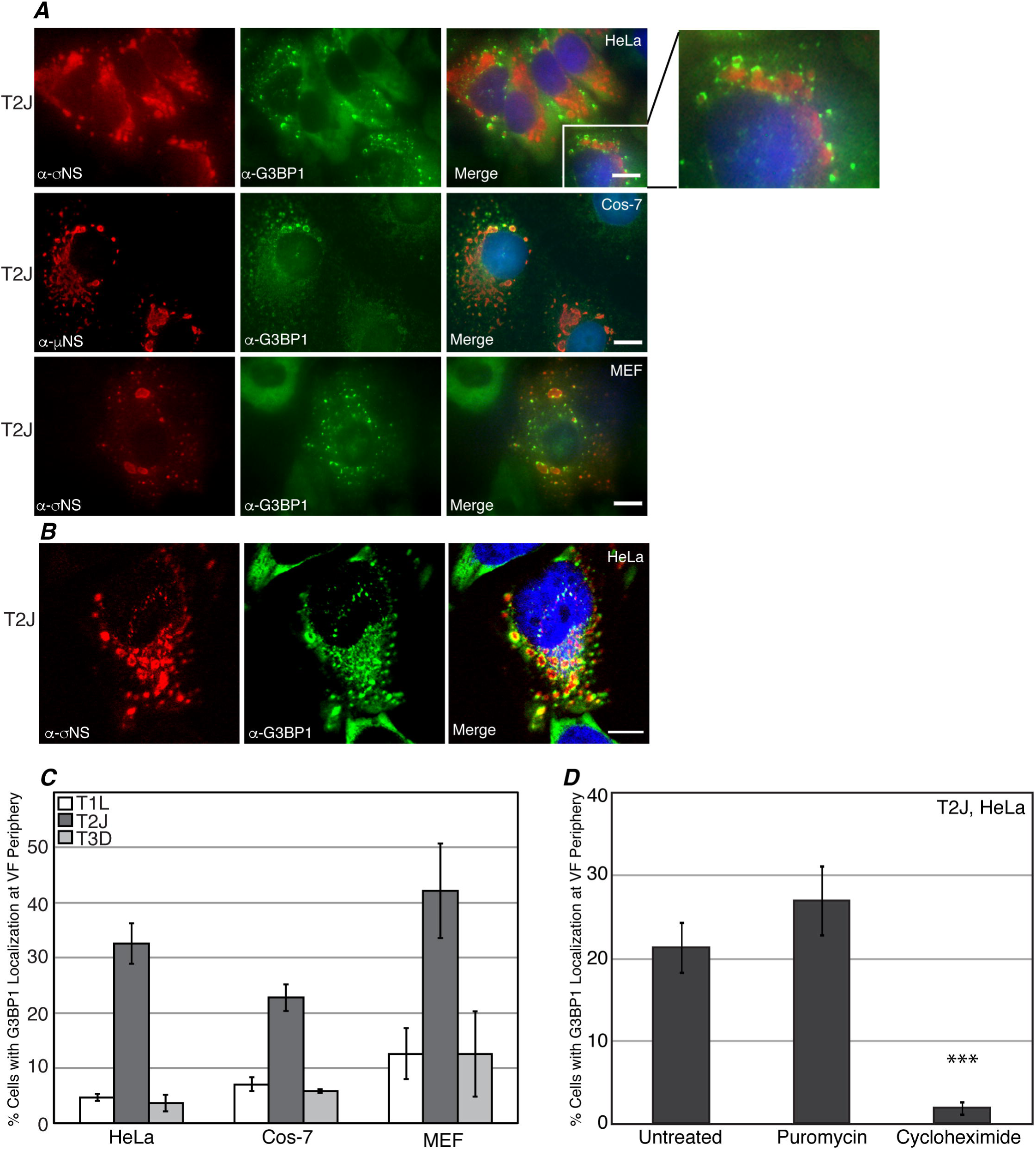
G3BP1 localizes to the outer periphery of VFs in MRV infected cells. (A) HeLa (first row), Cos-7 (second row) and MEF (third row) cells were infected with MRV strain T2J. At 24 h p.i., cells were fixed and immunostained with mouse α-σNS monoclonal (HeLa and MEF) or mouse α-μNS polyclonal (Cos-7) antibody (left columns) and rabbit α-G3BP1 polyclonal antibody (middle columns), followed by Alexa 594-conjugated donkey α-mouse IgG and Alexa 488-conjugated donkey α-rabbit IgG. Merged images containing DAPI-stained nuclei (blue) are shown (right columns). Bars=10 μm. The boxed inset (first row, third column) shows a magnified section of T2J infected HeLa cells. (B) HeLa cells were infected with MRV strain T2J. At 24 h p.i., the cells were fixed and immunostained with mouse α-σNS monoclonal antibody (left column) and rabbit α-G3BP1 polyclonal antibody (middle column), followed by Alexa 594-conjugated donkey α-mouse IgG and Alexa 488-conjugated donkey α-rabbit IgG and confocal images, including a merged image containing DAPI-stained nuclei (blue) are shown (right columns). Bars=10 μm. (C) HeLa, Cos-7 and MEF cells were infected with T1L (white bars), T2J (dark grey bars) or T3D (light grey bars), fixed and immunostained with mouse α-σNS monoclonal antibody and rabbit α-G3BP1 polyclonal antibody, followed by Alexa 594-conjugated donkey α-mouse IgG and Alexa 488-conjugated donkey α-rabbit IgG. Infected cells exhibiting G3BP1 localization at the VF periphery, out of the total number of infected cells, were counted and the means and standard deviations of three experimental replicates are shown. (D) HeLa cells were infected with MRV strain T2J. At 24 h p.i., cells were either untreated or treated with puromycin or cycloheximide for 1 h, fixed and immunostained with mouse α-σNS monoclonal antibody and rabbit α-G3BP1 polyclonal antibody followed by Alexa 594-conjugated donkey α-mouse IgG and Alexa 488-conjugated donkey α-rabbit IgG. Infected cells exhibiting G3BP1 localization at the VF periphery, out of the total number of infected cells, were counted and the means and standard deviations of three experimental replicates are shown.

The association of mRNA with polysomes is known to impact SG formation, with an influx of non-polysome associated mRNA promoting formation of SGs and an increase in polysome associated mRNA leading to inhibition of SG formation (46, 47). To determine whether the localization of G3BP1 around VFs during MRV infection is dependent on the state of mRNA translation in the cell, and therefore related to the induction of SG formation, we examined the percent of infected cells with G3BP1 localization to the VF periphery in the presence of puromycin, which dissociates polysomes, stimulating SG assembly (48), and cycloheximide, which prevents polysome dissociation, inhibiting SG assembly (47). HeLa cells were infected with T2J, and at 24 h p.i., cells were either left untreated, or treated with puromycin or cycloheximide for 1 h prior to fixation and immunostaining. Cells were then examined by immunofluorescence microscopy, and the number of infected cells with G3BP1 localization around VFs were counted (Fig. 1D). We found that the addition of puromycin did not lead to a significant increase in the number of infected cells with G3BP1 localization at the VF periphery. As puromycin promotes SG formation but does not independently induce SGs (46, 48), these results suggest that polysomal dissociation triggered by puromycin is not sufficient to drive G3BP1 localization around VFs in MRV infected cells that do not already exhibit the localization. Interestingly, however, the addition of cycloheximide led to a significant decrease in the number of infected cells with localization of G3BP1 around VFs, suggesting that inhibiting polysome dissociation prevents G3BP1 peripheral VF localization, and indicating that the altered localization of G3BP1 to VFs instead of canonical SGs is likely a result of active SG modulation during MRV infection.

The cycloheximide-dependent localization of G3BP1 around MRV VFs strongly suggests that there may be an association between MRV factories and cellular SGs. To further examine this, we investigated the localization of additional SG associated proteins in MRV infected cells. HeLa cells were infected with MRV strain T2J and at 24 h. p. i., cells were fixed and stained with antibodies against the MRV protein σNS, and SG-associated proteins Caprin1, USP10, TIA-1, TIAR or eIF3b followed by Alexa fluor-conjugated secondary antibodies and then examined by fluorescence microscopy (Fig. 2A, Fig. 2B). Our data showed that similar to G3BP1, Caprin1 (Fig. 2A, first row), USP10 (Fig. 2A, second row), TIAR (Fig. 2A, third row), TIA-1 (Fig. 2A, fourth row) or eIF3b (Fig. 2A, fifth row) localized to the outer periphery of VFs formed by MRV strain T2J in a subset of MRV infected cells (Fig. 2B). Taken together with our prior studies, these findings indicate that during later stages of MRV infection, rather than forming canonical SGs in response to phosphorylation of eIF2α induced by the virus or exogenous stressors such as sodium arsenite, SG associated proteins are either diffusely distributed as we previously reported (28, 36) or accumulate around MRV-induced VFs. Taken all together, these data suggest that VFs may play a role in SG modulation during MRV infection.

**Fig. 2.**
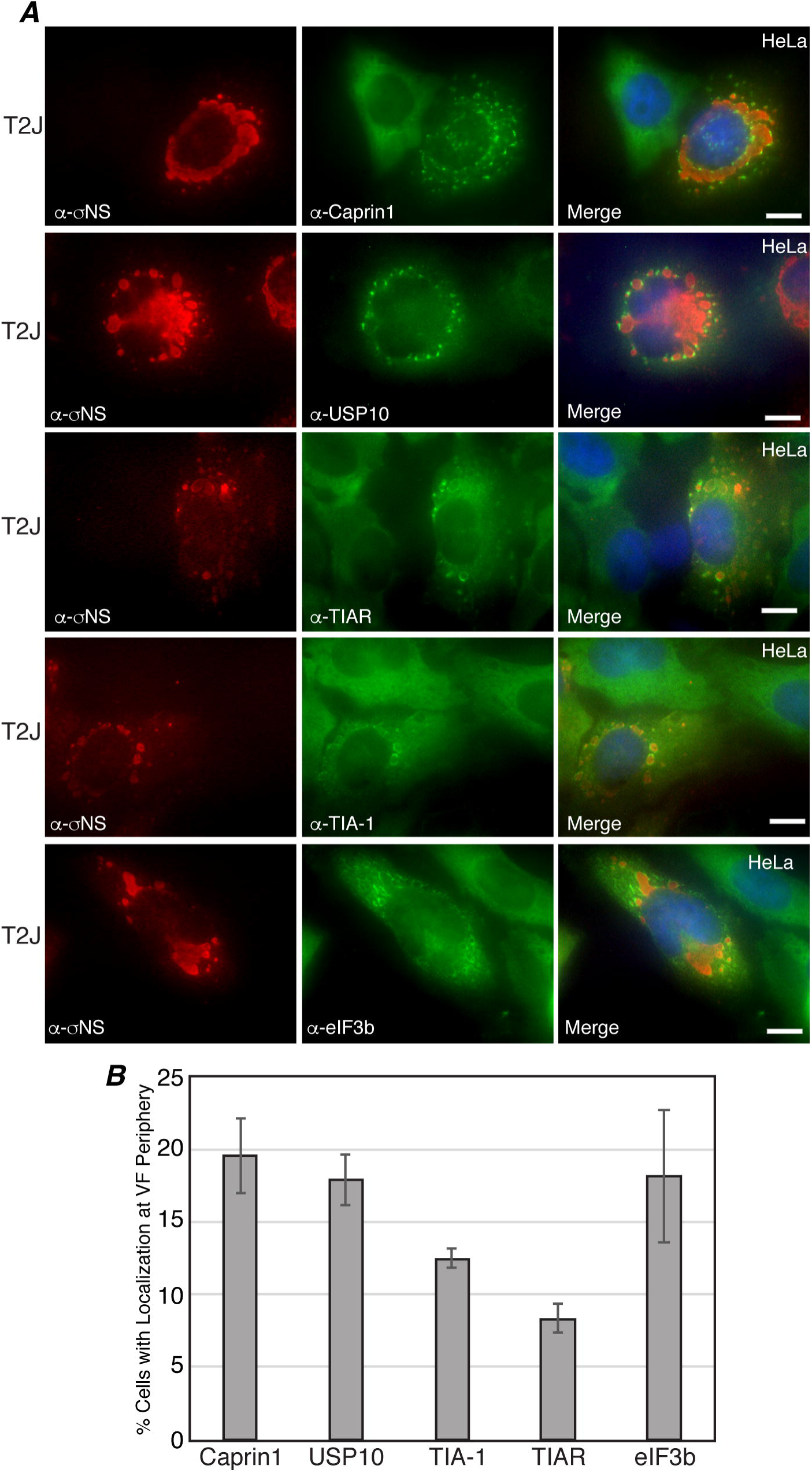
Multiple SG associated proteins localize to the VF periphery. (A) HeLa cells were infected with MRV strain T2J. At 24 h p.i., cells were fixed and immunostained with mouse α-σNS monoclonal antibody (left columns) and rabbit α-Caprin1 polyclonal antibody (middle column, first row), rabbit α-USP10 monoclonal antibody (middle column, second row), goat α-TIAR polyclonal antibody (middle column, third row), goat α-TIA-1 polyclonal antibody (middle column, fourth row) and rabbit α-eIF3b polyclonal antibody (middle column, fifth row) followed by Alexa 594-conjugated donkey α-mouse IgG (for σNS) and Alexa 488-conjugated donkey α-rabbit IgG (for Caprin1, USP10 and eIF3b) or Alexa 488-conjugated donkey α-goat IgG (for TIA-1, TIAR). Merged images containing DAPI-stained nuclei (blue) are shown (right columns). Bars=10 μm. (B) HeLa cells were infected with T2J, and fixed and immunostained as in A. Infected cells exhibiting Caprin1, USP10, TIA-1, TIAR and eIF3b localization at the VF periphery, out of the total number of infected cells, were counted and the means and standard deviations of three experimental replicates are shown.

### MRV σNS protein associates with G3BP1

When expressed alone in transfected cells, the MRV non-structural protein μNS forms structures that are strikingly similar to VFs (termed viral factory-like structures or VFLs), to which 6 other viral proteins and viral core particles are localized (25, 31, 41). Since μNS is involved in the nucleation of VFs and we have previously shown that μNS is localized to SA-induced stress granules (25, 31, 36, 41), we first investigated whether G3BP1 would be localized to VFLs formed by μNS expression in transfected cells. HeLa cells were transfected with plasmids encoding μNS, and 24 hours p. t., cells were fixed and stained with antibodies against μNS and G3BP1 followed by Alexa fluor-conjugated secondary antibodies to detect cellular localization of G3BP1 with respect to μNS-formed VFLs (Fig. 3A). G3BP1 was diffusely distributed throughout μNS transfected cells suggesting that expression of μNS alone is not sufficient to localize G3BP1 around VFLs (Fig. 3A, first row). The single-stranded RNA-binding non-structural protein σNS, exhibits a diffuse distribution throughout the cytoplasm and nucleus of infected cells when expressed alone, but is localized to VFLs when co-expressed with μNS (30, 32). Moreover, both σNS and G3BP1 associate with ribosomal proteins, suggesting they may have shared binding partners (10, 29). Both σNS and G3BP1 are diffusely distributed in cells, making it difficult to determine if the two proteins associate using a colocalization approach. (Fig. 3A, second row). However, overexpression of G3BP1 triggers SG assembly (7), therefore we induced SG formation in HeLa cells by co-transfecting a G3BP1 expressing plasmid with a plasmid expressing σNS. At 24 h p. t., cells were fixed and stained, and σNS and G3BP were visualized by immunofluorescence microscopy. In these experiments, G3BP1 formed SGs in many cells, and in some of those cells, σNS co-localized with G3BP1-induced SGs, suggesting that G3BP1 may associate with σNS (Fig. 3A, third row).

**Fig. 3.**
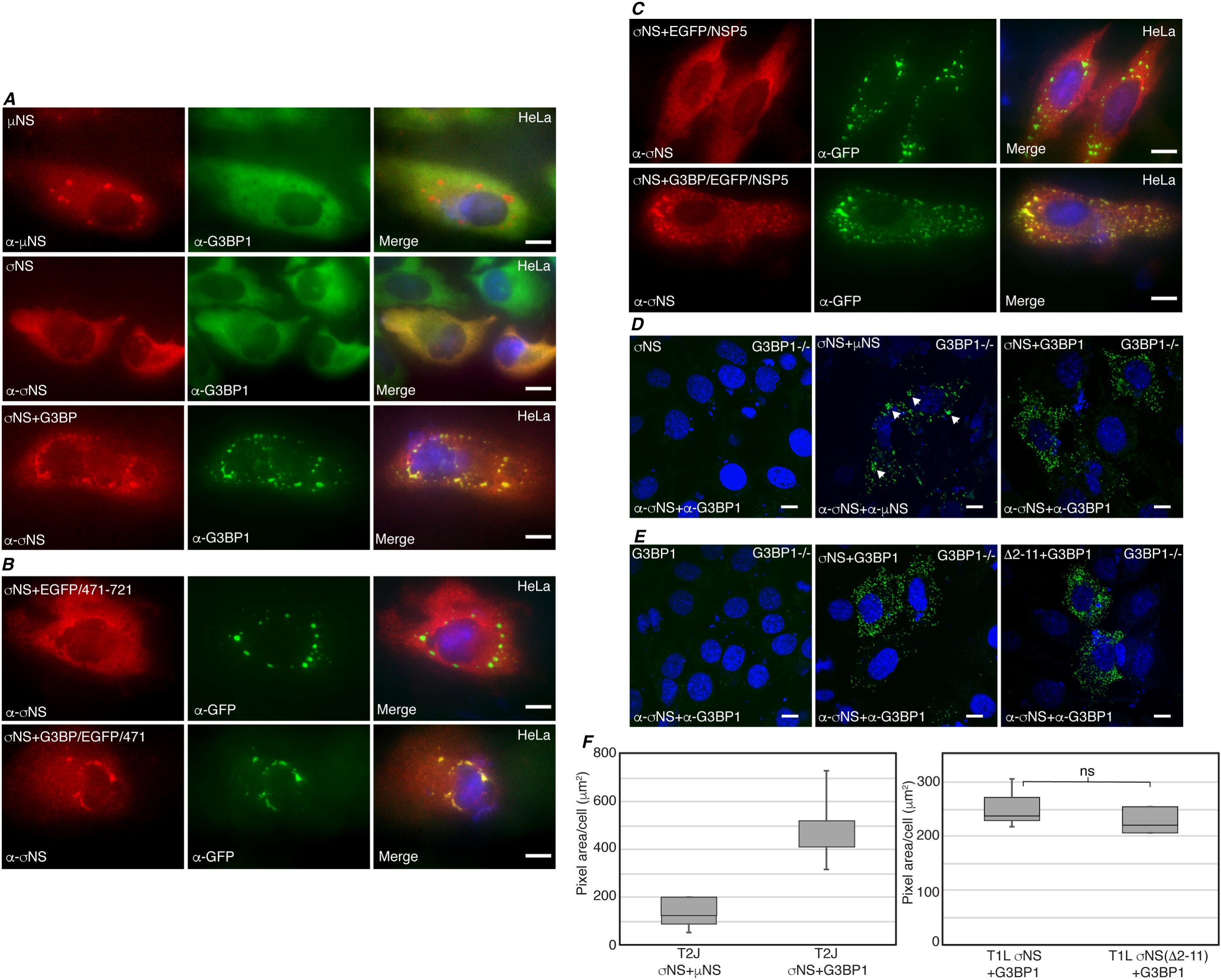
σNS associates with G3BP1. (A) HeLa cells were transfected with pCI-μNS (T1L) (first row), pCI-σNS (T1L) (second row) or pCI-σNS (T1L) and pCI-G3BP1 (third row). At 24 h p. t., cells were fixed and immunostained with mouse α-μNS polyclonal antiserum (left column, first row), or mouse α-σNS monoclonal antibody (left column, second and third rows) and rabbit α-G3BP1 polyclonal antibody (middle column, all rows), followed by Alexa 594-conjugated donkey α-mouse IgG and Alexa 488-conjugated donkey α-rabbit IgG. Merged images containing DAPI-stained nuclei (blue) are shown (right columns). Bars=10 μm. (B) HeLa cells were transfected with pCI-σNS and p-EGFP/μNS(471-721) (first row), or pCI-σNS and p-G3BP1/EGFP/μNS(471-721) (second row). At 24 h p. t., cells were fixed and immunostained with mouse α-σNS monoclonal antibody (left columns), and rabbit α-EGFP polyclonal antibody (middle columns), followed by Alexa 594-conjugated donkey α-mouse IgG and Alexa 488-conjugated donkey α-rabbit IgG. Merged images containing DAPI-stained nuclei (blue) are shown (right columns). Bars=10 μm. (C) HeLa cells were transfected with pCI-σNS and p-EGFP/NSP5 (first row), or pCI-σNS and p-G3BP1/EGFP/NSP5 (second row). At 24 h p. t., cells were fixed and immunostained with mouse α-σNS monoclonal antibody (left columns), and rabbit α-EGFP polyclonal antibody (middle columns), followed by Alexa 594-conjugated donkey α-mouse IgG and Alexa 488-conjugated donkey α-rabbit IgG. Merged images containing DAPI-stained nuclei (blue) are shown (right columns). Bars=10 μm. (D) G3BP1 -/- MEF cells were transfected with pCI-σNS (T2J, left column), pCI-σNS and pCI-μNS (T2J, middle column), or pCI-σNS (T2J) and pCI-G3BP1 (right column) and at 24 h p. t., subjected to PLA using mouse α-σNS monoclonal antibody (all columns), rabbit α-μNS polyclonal antibodies (middle column) or rabbit α-G3BP1 polyclonal antibodies (left and right columns) followed by mouse minus and rabbit plus antibody probes, ligation and amplification reactions. Merged confocal images containing DAPI-stained nuclei are shown. Bars=10 μm. (E) G3BP1-/- MEF cells were transfected with pCI-G3BP1 (all columns), pCI-σNS (T1L, middle column), or pCI-σNS(Δ2-11) (T1L, right column), and at 24 h p. t., subjected to PLA using mouse α-σNS monoclonal antibodies and rabbit α-G3BP1 polyclonal antibodies followed by mouse minus and rabbit plus antibody probes, ligation and amplification reactions. Merged confocal images containing DAPI-stained nuclei are shown. Bars=10 μm. (F) Images from D and E were subjected to 3D Objects Counter analysis (ImageJ) and pixel area/cell for each experimental condition was calculated and is shown. ns, not significant.

To further explore the association of σNS and G3BP1, we took advantage of published data showing that the carboxyl terminal one-third of μNS, spanning residues 471-721, is sufficient for formation of VFLs in transfected cells (41) but is unable to associate with σNS since it lacks the amino terminal residues 1-13 required for σNS binding (32). We designed a plasmid expressing a fusion protein, G3BP1/EGFP/μNS(471-721), by incorporating G3BP1 on the amino terminus of EGFP fused to the μNS (471-721) protein fragment. HeLa cells were transfected with pCI-σNS and either p-EGFP/μNS(471-721) (Fig. 3B, first row) or p-G3BP1/EGFP/μNS(471-721) (Fig. 3B, second row). Cells were fixed and stained with antibodies against σNS and EGFP, followed by Alexa fluor-conjugated secondary antibodies. While σNS did not associate with the structures formed by the EGFP/μNS(471-721) protein fusion, σNS did co-localize with structures formed by G3BP1/EGFP/μNS(471-721) in some cells, suggesting that VFLs formed by μNS(471-721) provide a platform for presenting G3BP1 to associate with σNS, independent of the association between σNS and μNS.

Because the μNS(471-721) protein fragment provided a foundation for presenting G3BP1 to σNS in the above experiments, there remained a possibility that this portion of μNS was contributing to G3BP1 association with σNS. To rule out any role of μNS in G3BP1 association with σNS, we utilized the rotavirus non-structural protein NSP5, which is functionally similar to μNS in terms of forming rotavirus viroplasm-like structures, without sharing substantial sequence homology with μNS (49, 50). We designed a plasmid construct in which the G3BP1 gene was fused to the amino terminus of an EGFP/NSP5 fusion plasmid, and transfected HeLa cells with pCI-σNS and p-EGFP/NSP5 (Fig. 3C, first row) or p-G3BP1/EGFP/NSP5 (Fig. 3C, second row). Cells were fixed and stained for σNS and EGFP, followed by Alexa fluor-conjugated secondary antibodies. Both EGFP/NSP5 and G3BP1/EGFP/NSP5 formed viroplasm-like stuctures, however, σNS only co-localized with structures formed by G3BP1/EGFP/NSP5 (Fig 3C, second row), and not with those formed by EGFP/NSP5 (Fig. 3C, first row). These data strongly suggest that G3BP1 association with σNS is not dependent on μNS.

To provide further evidence that G3BP1 associates with σNS, we performed a proximity ligation assay (PLA) (Fig. 3D and Fig. 3F). Plasmids expressing T2J σNS or G3BP1 alone (negative controls), T2J σNS and μNS (positive control), or G3BP1 and T2J σNS together were transfected into G3BP1 -/- MEF cells. At 18 h p.t., cells were fixed and stained with mouse antibodies against σNS and rabbit antibodies against μNS or G3BP1. Cells were incubated with plus and minus probe anti-rabbit and anti-mouse antibodies respectively, followed by ligation and amplification reactions. In this assay if the binding sites of the primary antibodies are within 30-40 nm, the ligation of oligos present in the ligation reaction will result in rolling circle replication during the amplification reaction, and fluorescent probes present in the amplification mix will bind and be apparent as fluorescent spots in locations within the cell where the interaction is taking place. Interestingly, as σNS and μNS associate primarily within VFLs, we routinely detected small dots of PLA amplification within larger spots that were presumably VFLs in our positive control samples (Fig. 3D, middle panel, arrowheads). Importantly, we detected abundant bright fluorescent spots in cells containing both σNS and G3BP1, but not in negative control cells missing either one of the two proteins (Fig. 3D, first panel, Fig. 3E, first panel), indicating that the two proteins are within 30-40 nm of each other, and further supporting our findings that σNS and G3BP1 associate within the cell. Finally, as both σNS and G3BP1 are single-stranded RNA binding proteins (5, 51), we wanted to determine if the σNS association with G3BP1 was occurring through σNS RNA binding. It has previously been shown that the σNS amino terminal residues 2-11 are necessary for RNA binding (52). We transfected G3BP1 -/- MEFs with plasmids expressing G3BP1 and either wildtype T1L σNS, or T1L σNS with a deletion of amino-acids 2-11 (σNSΔ2-11), and performed PLA (Fig. 3E). We found that wildtype T1L σNS, similar to T2J σNS strongly associated with G3BP1, suggesting the interaction is not strain-specific. Moreover, we found that deletion of σNS amino-acids 2-11 had no significant impact on the association, suggesting that σNS binding to RNA is not necessary for the association (Fig. 3E, third panel, Fig. 3F). Taken all together, these findings strongly suggest that G3BP1 associates with σNS and that this association is independent of viral strain, σNS RNA-binding, or σNS-μNS association.

### Expression of μNS and σNS is essential for G3BP1 localization around VFLs

Our results indicated that G3BP1 is recruited to the outer periphery of VFs in MRV infected cells, and in transfected cells, G3BP1 is not recruited to VFLs formed by μNS alone but interacts with σNS, a protein that is localized to VFLs by association with μNS (28). Based on these findings, we hypothesized that G3BP1 may be recruited to the outer periphery of VFLs formed by μNS via its association with σNS. To test our hypothesis, we transfected HeLa cells with plasmids encoding the μNS and σNS proteins from MRV strains T1L, T2J and T3D. At 24 h p.t., cells were fixed and stained with antibodies against σNS and G3BP1 followed by Alexa fluor-conjugated secondary antibodies, and the localization of σNS and endogenous G3BP1 was examined by immunofluorescence microscopy. As has been previously shown, σNS strongly co-localized with VFLs formed by μNS in these experiments. Confirming our hypothesis, we found that G3BP1 localization was visible as discrete staining around the outer margins of the VFLs formed by μNS derived from T1L, T2J and T3D viruses (Fig. 4A-first, second and third rows, respectively). Moreover, unlike what we measured in infected cells, there was no strain-specific difference in the percentage of cells (between 40-50% of all transfected cells) that exhibited G3BP1 localization to VFLs formed by μNS and σNS (Fig 4., and data not shown) suggesting that the MRV strain-specific differences we measured in infected cells were not a consequence of differences in σNS binding to G3BP1 or σNS recruitment of G3BP1 to VFLs.

**Fig. 4.**
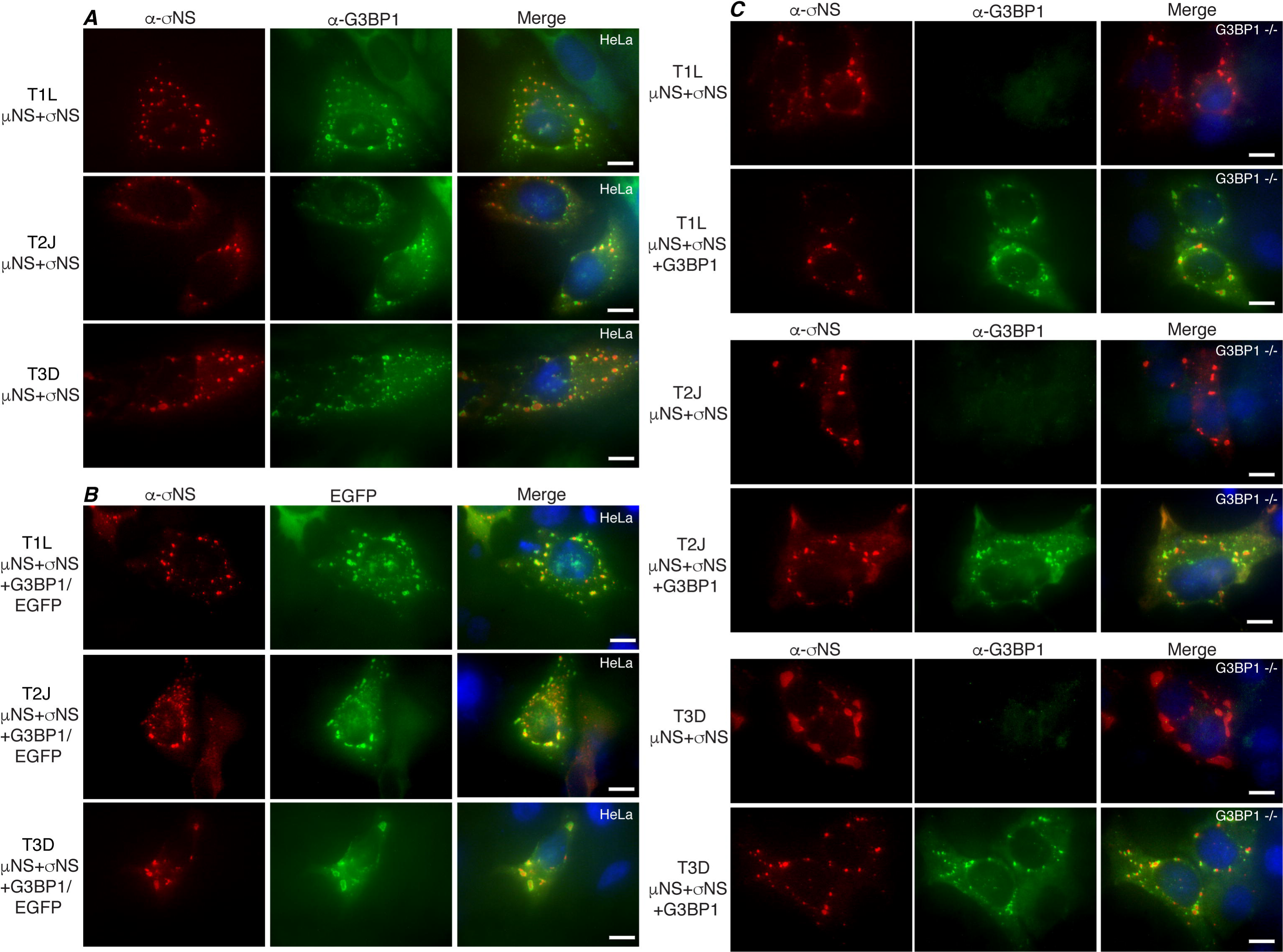
G3BP1 is recruited to the periphery of VFLs by σNS and μNS. (A) HeLa cells were transfected with pCI-μNS and pCI-σNS of T1L (first row), T2J (second row), or T3D (third row). At 24 h p. t., cells were fixed and immunostained with mouse α-σNS monoclonal antibody (left columns) and rabbit α-G3BP1 polyclonal antibody (middle columns), followed by Alexa 594-conjugated donkey α-mouse IgG and Alexa 488-conjugated donkey α-rabbit IgG. Merged images containing DAPI-stained nuclei (blue) are shown (right columns). Bars=10 μm. (B) HeLa cells were transfected with pCI-μNS and pCI-σNS of T1L (first row), T2J (second row), or T3D (third row) and p-G3BP1/EGFP. At 24 h p. t., cells were fixed and immunostained with mouse α-σNS monoclonal antibody (left columns) followed by Alexa 594-conjugated donkey α-mouse IgG. The inherent fluorescence of EGFP was used to detect G3BP1/EGFP (middle columns). Merged images containing DAPI-stained nuclei (blue) are shown (right columns). Bars=10 μm. (C) G3BP1 -/- MEF cells were transfected with pCI-μNS and pCI-σNS of T1L (top two panels), T2J (middle two panels), or T3D (bottom two panels) with pCI-G3BP1 (bottom rows) or without pCI-G3BP1 (top rows). At 24 h p. t., the cells were fixed and immunostained with mouse α-σNS monoclonal antibody (left columns) and rabbit α-G3BP1 polyclonal antibody (middle columns), followed by Alexa 594-conjugated donkey α-mouse IgG and Alexa 488-conjugated donkey α-rabbit IgG. Merged images containing DAPI-stained nuclei (blue) are shown (right columns). Bars=10 μm.

In order to confirm the recruitment of G3BP1 to the outer periphery of VFLs, and to rule out possible cross-reaction of the G3BP1 antibody, we constructed a plasmid in which the G3BP1 protein was fused to the amino terminus of EGFP. G3BP1 localization was then monitored by examining the inherent fluorescence of the G3BP1/EGFP fusion protein. HeLa cells were co-transfected with plasmids expressing μNS and σNS of either T1L (Fig. 4B, first row) T2J (Fig. 4B, second row) or T3D (Fig. 4B, third row) and p-G3BP1/EGFP. Similar to endogenous G3BP1, G3BP1/EGFP was strongly localized to the outer periphery of VFLs formed by σNS and μNS derived from all three virus strains in transfected cells, indicating that this phenotype is not limited to endogenous G3BP1 or an artifact of the G3BP1 antibody.

We further examined this phenotype by co-expressing σNS and μNS of T1L (Fig. 4C, first panel), T2J (Fig. 4C, second panel), or T3D (Fig. 4C, third panel) in G3BP1 -/- MEFs with (bottom rows) or without (top rows) plasmid expressed G3BP1, and monitored the localization of G3BP1 with respect to σNS. Similar to our previous results, G3BP1 displayed a peripheral localization along the outside margins of the VFLs formed by σNS and μNS from all three MRV strains further confirming this phenotype. Taken together, these results indicate that the MRV non-structural protein σNS interacts with endogenous or exogenously expressed G3BP1 around the margins of VFLs formed by μNS, suggesting that co-expression of μNS and σNS is essential for G3BP1 localization to the outer peripheries of VFs in infected cells. Since there was no obvious strain specific difference in G3BP1 localization around σNS/μNS formed VFLs, we continued our investigation with σNS and μNS derived from T2J for the following experiments.

In infected cells, our data suggested that in addition to G3BP1, SG proteins Caprin1, TIA-1, TIAR and eIF3b were also all localized around VFs (Fig. 1). Because the expression of μNS and σNS was necessary for the recruitment of G3BP1 to VFLs in transfected cells (Fig. 4), we investigated the role of σNS and μNS co-expression in localization of the other SG proteins to VFLs (Fig. 5). HeLa cells were transfected with plasmids expressing σNS and μNS from MRV strain T2J, and immunostained with antibodies against σNS and Caprin1, TIAR, TIA-1 or eIF3b. As seen in infected cells, Caprin1 (Fig. 5, first row), TIAR (Fig. 5, second row), TIA-1 (Fig. 5, third row) and eIF3b (Fig. 5, fourth row) displayed a pattern of localization similar to G3BP1, characterized as distinct circling around the outer margins of the VFLs in many transfected cells. These results provide additional evidence that the MRV non-structural protein σNS and μNS are key players in driving the localization of SG-associated proteins around VFLs in transfected cells and VFs in infected cells.

**Fig. 5.**
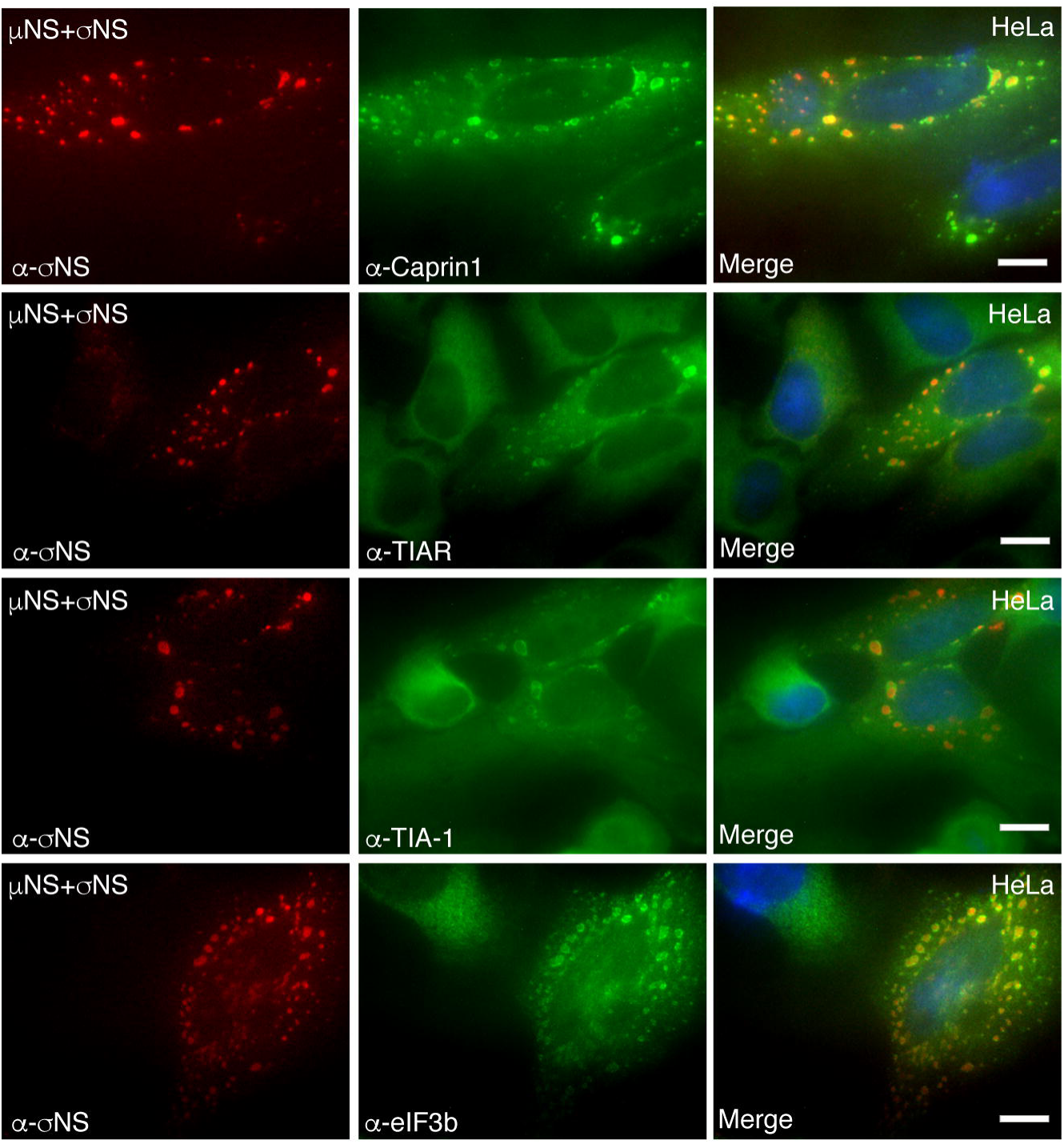
Multiple SG associated proteins are recruited to the periphery of VFLs by σNS and μNS. HeLa cells were transfected with pCI-μS (T2J) and pCI-σNS (T2J). At 24 h p. t., cells were fixed and immunostained with mouse α-σNS monoclonal antibody (left columns) and rabbit α-Caprin1 polyclonal antibody (middle column, first row), goat α-TIAR polyclonal antibody (middle column, second row), goat α-TIA-1 polyclonal antibody (middle column, third row), or rabbit α-eIF3b polyclonal antibody (middle column, fourth row) followed by Alexa 594-conjugated donkey α-mouse IgG and Alexa 488-conjugated donkey α-rabbit IgG or Alexa 488-conjugated donkey α-goat IgG. Merged images containing DAPI-stained nuclei (blue) are shown (right columns). Bars=10 μm.

### Localization of SG proteins to VFLs is G3BP1-dependent

Caprin1 has been shown to directly associate with G3BP1, while TIA-1, TIAR and eIF3b localize to SGs, but are not known to directly associate with G3BP1 (7). Because all these proteins localized around the peripheries of VFs in infected cells (Fig. 2) and VFLs in cells transfected with σNS and μNS (Fig. 5), via dual σNS association with μNS and G3BP1, we wanted to determine if the localization of these proteins to the VFL periphery was mediated by G3BP1. To address any potential impact of the G3BP1 homologue, G3BP2, on this localization we performed our experiments in cells lacking both G3BP1 and G3BP2. Wild-type and ΔΔG3BP1/2 U2OS cells were co-transfected with plasmids expressing T2J σNS and μNS. Cells were stained for σNS and either G3BP1 (Fig. 6A), Caprin1 (Fig. 6B), TIAR (Fig. 6C), TIA-1 (Fig. 6D) or eIF3b (Fig. 6E) followed by Alexa fluor-conjugated secondary antibodies, then visualized by immunofluorescence microscopy. Our results indicate that in the absence of G3BP, Caprin1, TIA-1 and TIAR were not localized to VFLs in any cells suggesting that the recruitment of these proteins to VFLs is G3BP dependent. However, unlike other SG proteins, eIF3b still exhibited VFL peripheral localization in many cells even in the absence of G3BP suggesting that eIF3b recruitment to VFL is independent of G3BP and is potentially due to the previously described presence of translational initiation complexes around VFs (29). Taken together these results suggest that the association of σNS with SG effector G3BP plays a significant role in driving the localization of other SG proteins to the VFL periphery. Moreover, as TIA-1/R do not directly associate with G3BP and simply co-localize with G3BP in SGs, this data also indicates that MRV factory formation by μNS, and σNS association with μNS and G3BP1 may result in the redistribution of SGs from their normal localization in the cell to the periphery of VFs.

**Fig. 6.**
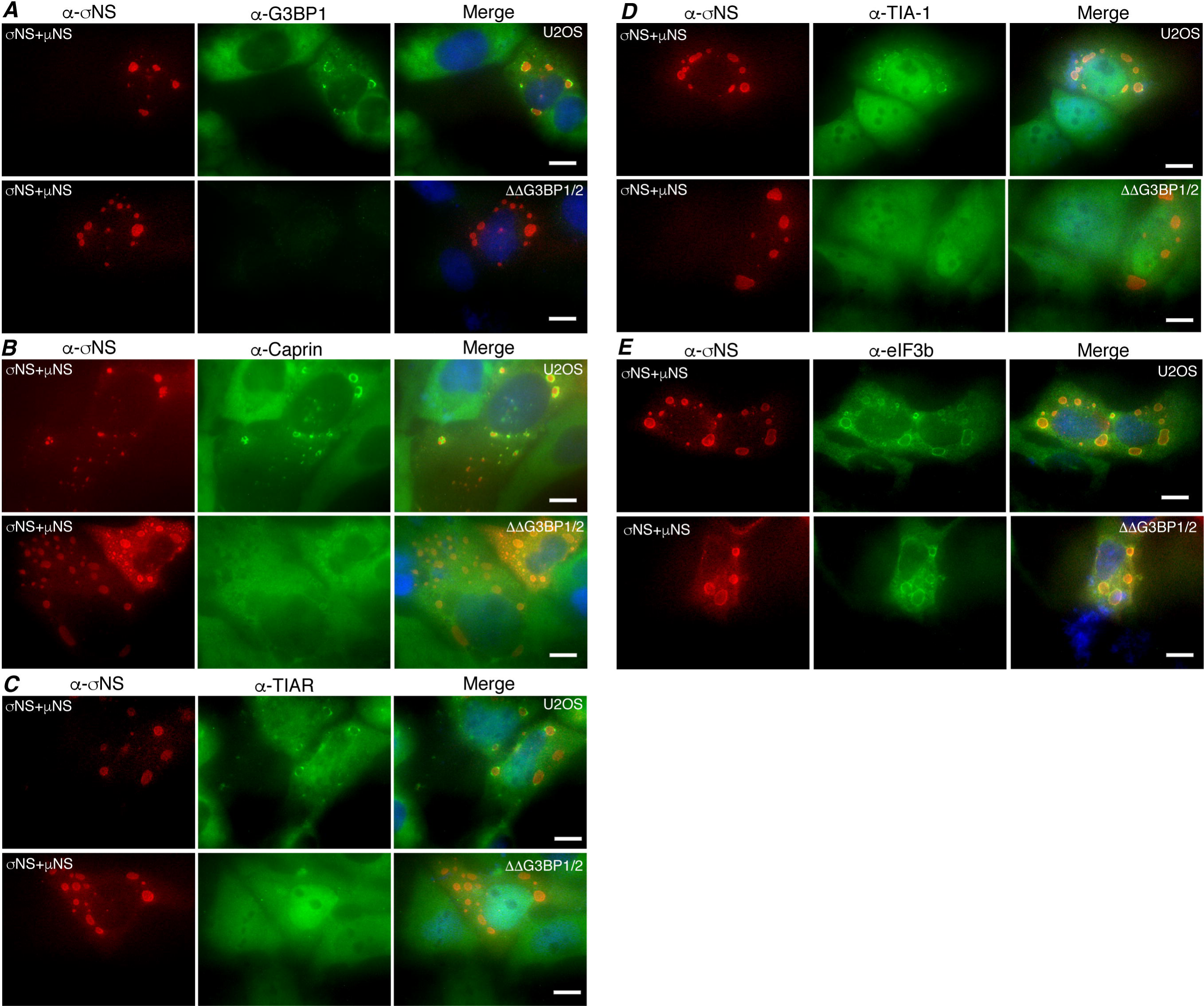
SG associated protein localization to VFLs is G3BP1 dependent. Wildtype (top rows) and ΔΔG3BP1/2 (bottom rows) U2OS cells were transfected with pCI-μNS (T2J) and pCI-σNS (T2J). At 24 h p. t., cells were fixed and immunostained with mouse α-σNS monoclonal antibody (left columns) and (A) rabbit α-G3BP1 polyclonal antibody, (B) rabbit α-Caprin1 polyclonal antibody, (C) goat α-TIAR polyclonal antibody (D) goat α-TIA-1 polyclonal antibody and (E) rabbit α-eIF3b polyclonal antibody followed by Alexa 594-conjugated donkey α-mouse IgG and Alexa 488-conjugated donkey α-rabbit or Alexa 488-conjugated donkey α-goat IgG. Merged images containing DAPI-stained nuclei (blue) are shown (right columns). Bars=10 μm.

### The C-terminal RGG and RRM domains of G3BP1 are involved in VFL localization and σNS interaction

The amino terminal NTF2-like domain of G3BP1 is the most highly conserved domain between species and within the mammalian G3BP1 family (53) and is essential for G3BP1 association with Caprin1 and USP10 (54). Site directed mutagenesis at residue 33 of G3BP1 from phenylalanine to tryptophan (G3BP1-F33W) disrupts the ability of G3BP1 to bind Caprin1 or USP10 (54) and reduces interaction with PKR (55), however this mutant remains SG competent (10). Phosphorylation of G3BP1 at residue 149 disrupts its ability to nucleate SGs (7), and, when expressed endogenously, the G3BP1 phosphomimetic mutant S149E is unable to form SGs (7) suggesting that residue 149 of G3BP1 functions as an SG regulatory switch (10). The carboxyl terminus of G3BP1 mediates G3BP1 binding to specific RNA sequences (56) and is involved in G3BP1 interaction with 40S ribosomal subunits (10). This region consists of predicted RNA binding domains, RRM (RNA recognition motif) and RGG (arginine-glycine rich) (57), which are also involved in G3BP1 mediated SG modulation (10). The RGG domain is required for SA induced SG formation whereas the RRM domain is not (9). As our findings indicated that G3BP is critical for localization of other SG associated proteins to VFLs, we sought to determine the specific site/domain that is involved in localization around VFLs. We created three amino terminus site specific mutants, G3BP1-F33W, G3BP1-S149A and G3BP1-S149E and three carboxyl terminus deletion mutants, G3BP1-ΔRRM (residues 340-416), G3BP1-ΔRGG (residues 428-456) and G3BP1-ΔRRM/ΔRGG in G3BP1 expression plasmids (Fig. 7A) and investigated which particular site or domain of G3BP1 is involved in localization of G3BP1 to VFLs. G3BP1 -/- MEF cells were transfected with plasmids expressing either wildtype or mutant G3BP1, with plasmids encoding T2J σNS and μNS. Cells were then fixed and stained with antibodies against G3BP1 and σNS, followed by Alexa fluor-conjugated secondary antibodies, and examined by immunofluorescence microscopy. G3BP1-F33W (Fig. 7B, second row), G3BP1-S149A (Fig. 7B, third row), and G3BP1-S149E (Fig. 7B, fourth row) were localized around VFLs in the presence of σNS/μNS expression similar to wildtype G3BP1 (Fig. 7B, first row), suggesting that Caprin1/USP10 binding, or modulation of phosphorylation of amino-acid 149 do not play a role in G3BP1 localization to VFLs. However, G3BP1-ΔRRM (Fig. 7B, fifth row), G3BP1-ΔRGG (Fig. 7B, sixth row) or G3BP1-ΔRRM/ΔRGG (Fig. 7B, seventh row) displayed significantly reduced VFL localization characterized by either weakened or absent G3BP1 staining around the VFL periphery in the presence of σNS/μNS expression. Quantification of these results by calculating the percentage of cells exhibiting any detectable G3BP1 localization around VFLs demonstrated that the RRM domain led to an average 30% reduction, deletion of the RGG domain led to an average 50% reduction, and deletion of both the RRM and RGG domains led to near complete disruption of G3BP1 localization to VFLs respectively (Fig. 7C).

**Fig. 7.**
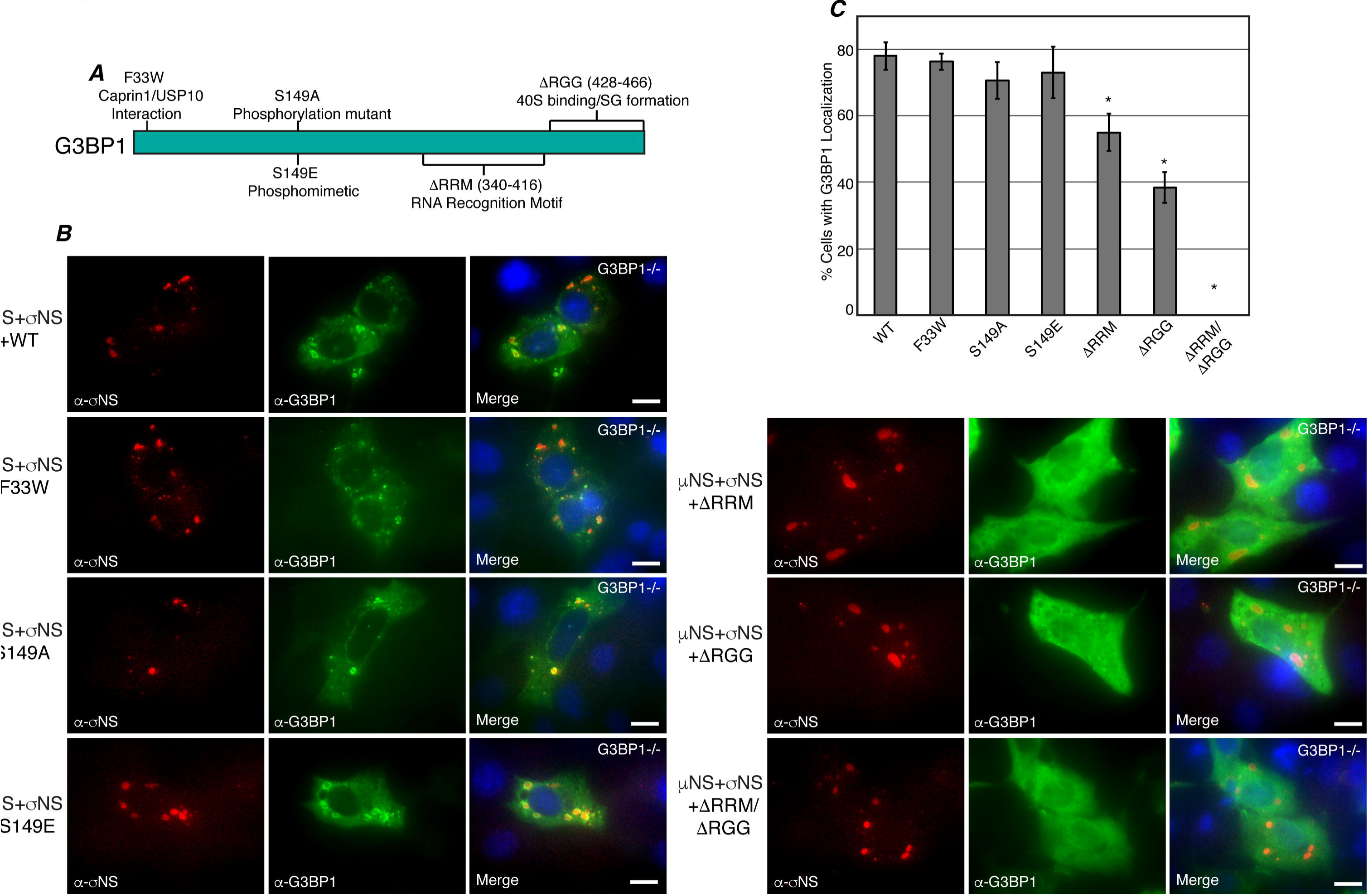
G3BP1 C-terminus is necessary for VFL localization. (A) Illustration of G3BP1 gene mutations used for mapping the region(s) necessary for G3BP1 recruitment to VFLs. (B) G3BP1 -/- MEF cells were transfected with pCI-μNS (T2J), pCI-σNS (T2J) and individual pCI-G3BP1 wildtype (first row) or mutant plasmids F33W (second row), S149A (third row), S149E (fourth row), ΔRRM (fifth row), ΔRGG (sixth row) or ΔRRMΔRGG (seventh row). At 24 h p. t., the cells were fixed and immunostained with mouse α-σNS monoclonal antibody (left columns) and rabbit α-G3BP1 polyclonal antibody (middle columns) followed by Alexa 594-conjugated donkey α-mouse IgG and Alexa 488-conjugated donkey α-rabbit IgG. Merged images containing DAPI-stained nuclei (blue) are shown (right columns). Bars=10 μm. (C) G3BP1 -/- MEF cells were treated as in (B). The percentage of G3BP1 -/- MEF cells exhibiting G3BP1 localization around VFLs out of the total number of cells were counted, and the means and standard deviations of three experimental replicates are shown. A statistically significant difference for which *p* < 0.05 is indicated with an asterisk (∗).

In order to determine if this loss of G3BP1 localization to VFLs correlated with σNS binding to G3BP1, we performed PLA with σNS and wildtype G3BP1, G3BP1-ΔRRM, G3BP1-ΔRGG, and G3BP1-ΔRRM/ΔRGG, in the absence and presence of μNS. G3BP1 -/- MEFs were transfected with plasmids expressing σNS, and each of the G3BP1 plasmids without and with a μNS expressing plasmid, and at 18 h p.t., cells were fixed, permeabilized, and subjected to PLA (Fig. 8A). Interestingly, in both the presence and absence of μNS, each tested G3BP1 mutant interaction with σNS was significantly diminished in both intensity and number of interaction sites in this assay (Fig. 8B). This suggests that the loss of G3BP1 localization to VFLs in these mutants correlates with a significant loss in G3BP1 binding to σNS, indicating that this interaction plays an important role in the re-localization of SG associated proteins from canonical SGs to the VFL periphery in transfected and infected cells. Importantly, however, because the interaction of σNS and G3BP1 was not completely disrupted in the ΔRGG mutant, which no longer associates with ribosomes, these findings also indicate that σNS association with G3BP1 is not occurring through both σNS and G3BP1 binding to ribosomes. Moreover, it has previously been reported that the ΔRRM mutant has increased binding to ribosomes relative to wildtype G3BP1 (10). Because we see a significant decrease in both the localization of G3BP1 to VFLs and σNS binding with this mutant, this further suggests that this interaction and phenotype are independent of ribosomal binding. Finally, because loss of the domain is not sufficient to fully prevent σNS binding, these results also suggest that RNA binding by the G3BP1 RRM domain may not be necessary for σNS association with G3BP1. Taken together these data suggest that the entire C-terminal domain of G3BP1 is essential for both maximal σNS binding and VFL localization.

**Fig. 8.**
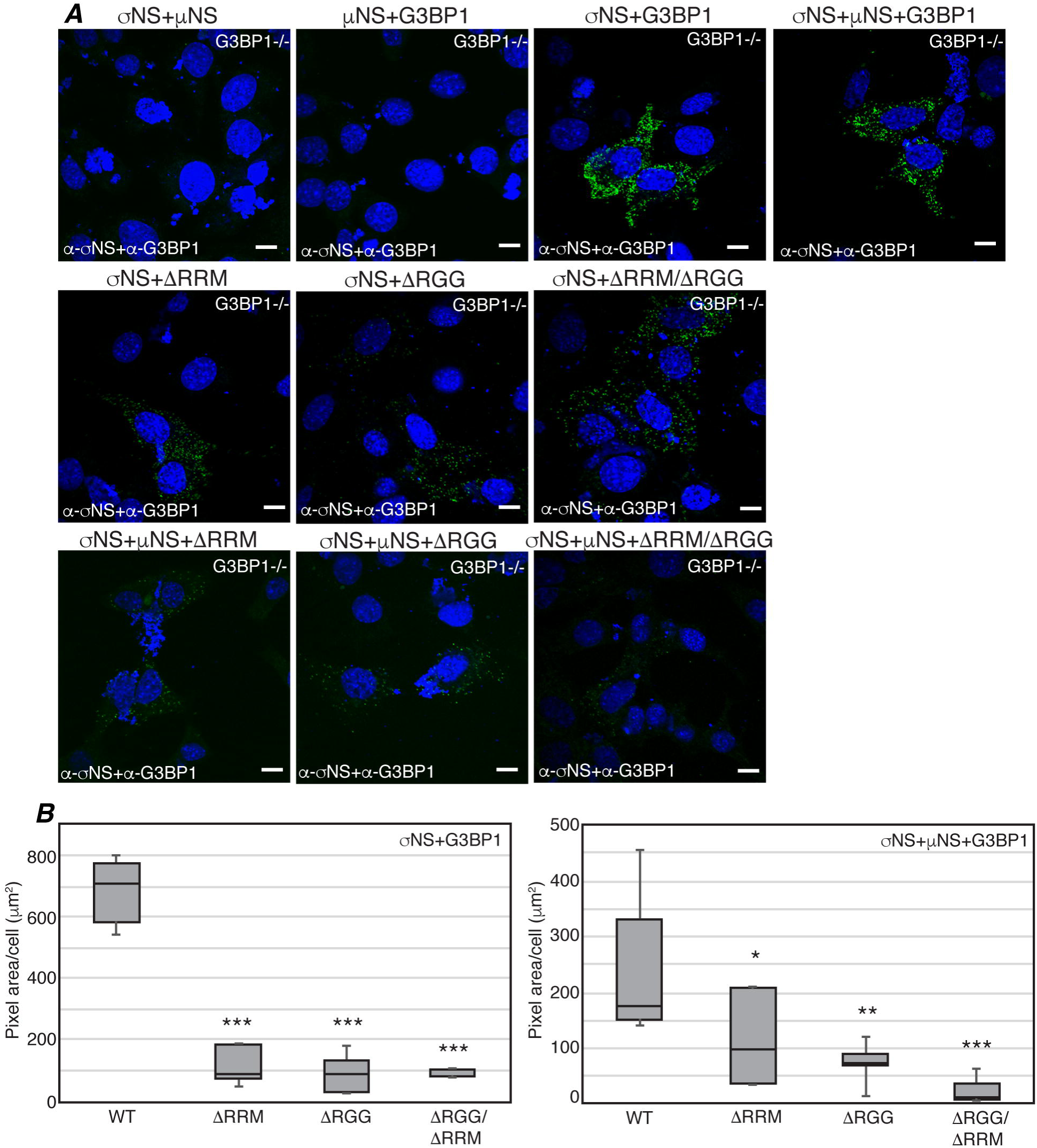
The G3BP1 C-terminal domains play a role in σNS binding. (A) G3BP1 -/- MEF cells were transfected with pCI-σNS (T2J, top row, first, third, and fourth columns; second row, all columns; third row, all columns), pCI-μNS (T2J, top row, second and fourth columns; third row, all columns), pCI-G3BP1 (top row, second, third, and fourth columns), G3BP1ΔRRM (second and third rows, first columns), G3BP1ΔRGG (second and third rows, second columns), or G3BP1 ΔRRM/ΔRGG (second and third rows, third columns) and at 24 h p.t., subjected to PLA using mouse α-σNS monoclonal antibody and rabbit α-G3BP1 polyclonal antibodies followed by mouse minus and rabbit plus antibody probes, ligation and amplification reactions. Merged confocal images containing DAPI-stained nuclei are shown. Bars=10 μm. (B) Images from A were subjected to 3D Objects Counter analysis (ImageJ) and pixel area/cell for each experimental condition was calculated and is shown. A statistically significant difference for which *p* < 0.05 or *p* < 0.01 or *p* < 0.001 were considered statistically significant and are indicated with one (^∗^) or two (^∗∗^) or three (^∗∗∗^) asterisks in the figures respectively.

### Co-expression of μNS and σNS interferes with SG formation

Previously we have reported that during later stages of infection with all three MRV strains (T1L, T2J and T3D), SG formation was disrupted in most cells even in the presence of cellular stressors such as sodium arsenite (35). We wanted to determine if expression of μNS or σNS or co-expression of μNS and σNS interfered with formation of canonical SGs that are induced by SA treatment (Fig. 9A). HeLa cells were transfected with plasmids expressing either T2J σNS (Fig. 9A, first row) or T2J μNS (Fig. 9A, second row) or co-transfected with T2J σNS and μNS (Fig. 9A, third row). 24 h p.t., cells were treated with SA and immunostained for G3BP1 and σNS or μNS. Our results demonstrated that expression of σNS or μNS alone did not interfere with formation of canonical, SA-induced SGs as evident by G3BP1 staining in cells expressing either μNS or σNS (Fig. 9 first and second rows). Instead, as we have reported here (Fig. 3) and elsewhere (36), a subset of σNS and μNS in cells co-localized to SGs. However, in cells that expressed both μNS and σNS, localization of G3BP-containing SG puncta was completely altered and G3BP1 was either concentrated around the VFL periphery (Fig. 9A, third row) or diffusely distributed throughout the cell, in some cases with weak peripheral VFL localization. High resolution confocal images confirm that instead of large granules localized throughout the cytoplasm as is seen in non-transfected cells (Fig. 9B, first row), G3BP1 localization was drastically altered following SA treatment when σNS and μNS are co-expressed in cells, with cells demonstrating G3BP1 either strongly localized to the σNS/μNS periphery (Fig. 9B, second row), or G3BP1 localized diffusely throughout the cytoplasm, in some cases with weak VFL peripheral staining (Fig. 9B, third row). As SA is predicted to induce phosphorylation of eIF2α and SG formation in all cells, these data strongly suggest that the presence of σNS/μNS VFLs in cells interferes with normal SG formation.

**Fig. 9.**
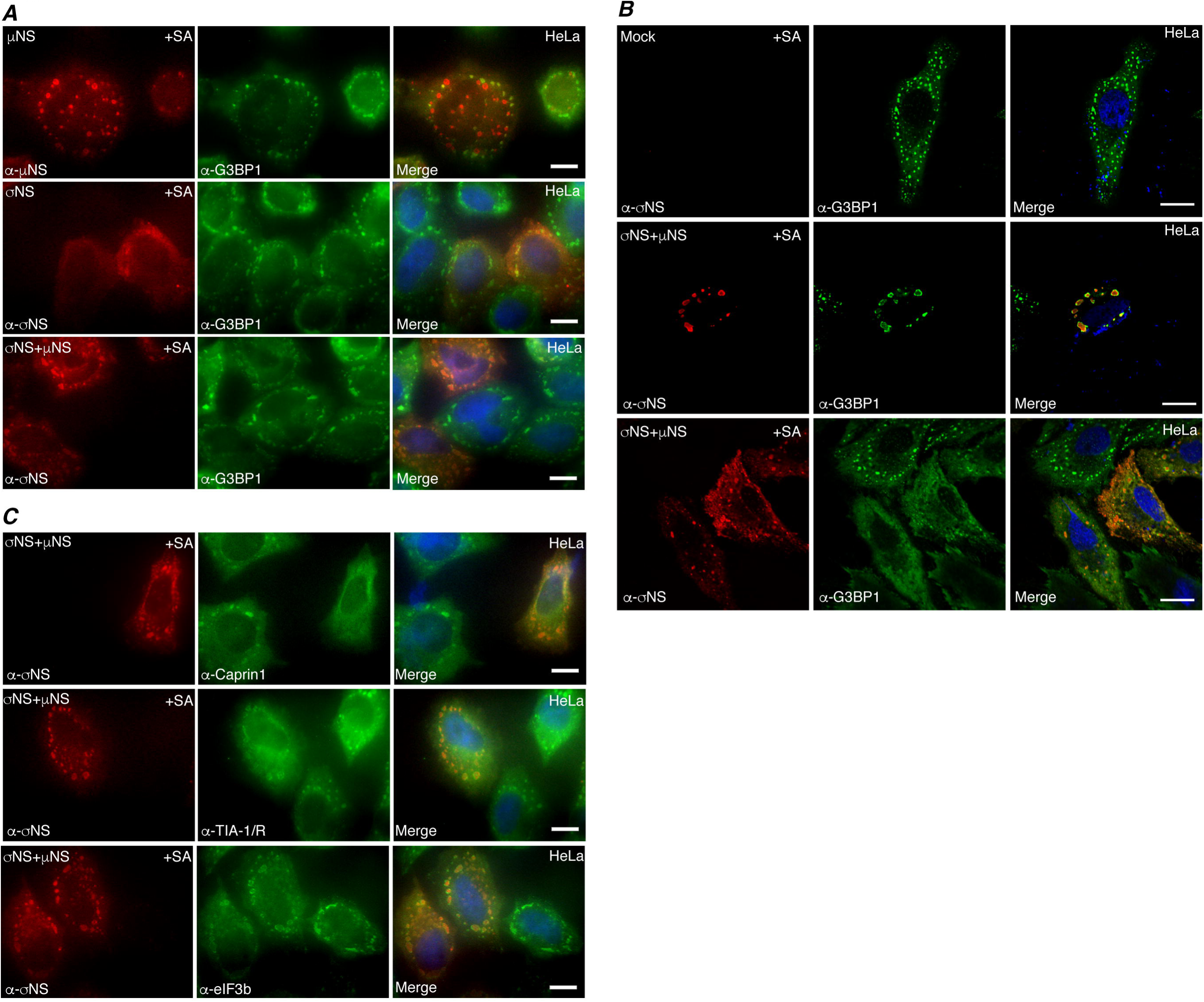
μNS and σNS expression interferes with canonical SG formation. (A) HeLa cells were transfected with pCI-μNS (T2J) (first row) or pCI-σNS (T2J) (second row) or pCI-σNS (T2J) and pCI-μNS (T2J) (third row). At 24 h p. t., cells were treated with SA, fixed and immunostained with mouse α-μNS polyclonal or mouse α-σNS monoclonal antibody (left columns) and rabbit α-G3BP1 polyclonal antibody (middle columns) followed by Alexa 594-conjugated donkey α-mouse IgG and Alexa 488-conjugated donkey α-rabbit IgG. Merged images containing DAPI-stained nuclei (blue) are shown (right columns). Bars=10 μm. (B) HeLa cells were mock-transfected (top row) or transfected with pCI-μNS (T2J) and pCI-σNS (T2J) (second and third rows). At 24 h p. t., cells were treated with SA for 45 min, fixed and immunostained with mouse α-σNS monoclonal antibody (left columns) and rabbit α-G3BP1 polyclonal antibody (middle columns) followed by Alexa 594-conjugated donkey α-mouse IgG and Alexa 488-conjugated donkey α-rabbit IgG. Merged images containing DAPI-stained nuclei (blue) are shown (right columns). Bars=10 μm. (C) HeLa cells were transfected with pCI-μNS (T2J) and pCI-σNS (T2J). At 24 h p. t., cells were treated with SA for 45 min, fixed and immunostained with mouse α-σNS monoclonal antibody (left columns) and rabbit α-Caprin1 polyclonal antibody (middle column, first row), goat α-TIA-1/R polyclonal antibody (middle column, second row) or rabbit α-eIF3b polyclonal antibody (middle column, third row) followed by Alexa 594-conjugated donkey α-mouse IgG and Alexa 488-conjugated donkey α-rabbit IgG or Alexa 488-conjugated donkey α-goat IgG. Merged images containing DAPI-stained nuclei (blue) are shown (right columns). Bars=10 μm.

Next we wanted to confirm whether the altered localization of G3BP1 in cells expressing σNS and μNS was impacting other SG associated proteins following SA treatment. HeLa cells were transfected with plasmids expressing T2J σNS and μNS and 24 h p.t., cells were treated with SA, then stained for σNS and major SG marker proteins, Caprin1 (Fig. 9C, first row), TIA-1/R (Fig. 9C, second row) and eIF3b (Fig. 9C, third row), followed by Alexa fluor-conjugated secondary antibodies, and visualized by immunofluorescence microscopy. Our results indicated that canonical SG proteins, Caprin1, TIA-1/R and eIF3b were also distributed either around VFL structures or diffusely throughout the cell, suggesting that co-expression of μNS and σNS leads to a striking change in distribution of these representative SG associated proteins away from canonical SA-induced SG puncta. Taken together, our results provide strong evidence that in cells where σNS and μNS are co-expressed, σNS binds G3BP1 and μNS to re-localize SG associated proteins in cells and interfere with normal formation of SA-induced SGs, suggesting a novel mechanism for SG disruption during MRV infection.

### G3BP1 inhibits MRV growth

Our data supports a hypothesis where σNS association with G3BP1 or a protein complex containing G3BP1, and recruitment of this complex to VFs may interfere with normal SG formation to prevent an inhibitory effect on infection. Therefore, we next wanted to examine the impact of G3BP1 on MRV replication. Wildtype and G3BP1-/- MEFs were infected with MRV strains T1L, T2J or T3D and at 0, 24, 48, and 72 h p.i., cells were harvested and subjected to three freeze/thaw cycles. Cell lysates were then subjected to standard plaque assays on L929 cells to determine the virus titers in each cell type over time (Fig. 10). Deletion of G3BP1 facilitated increased T1L and T2J growth over time as indicated by a statistically significant higher virus titer respectively in the G3BP1 knockout MEFs at 24, 48, and 72 h p.i. relative to wildtype MEFs. However, G3BP1 deletion did not appear to have substantial impact on T3D growth as there was no significant difference in T3D virus titer even at 72 h p.i. To further confirm our findings and to rule out the possibility that in the absence of G3BP1, G3BP2 might play a role in regulating MRV replication, we performed replication assays in wildtype, ΔG3BP1 and ΔΔG3BP1/2 U2OS cells. Our results suggested that again in the absence of G3BP1 or in the absence of both G3BP1/2, replication of MRV strains T1L and T2J was significantly increased by 5-10 fold over a period of 24, 48, and 72 h p.i., while there was again no impact on T3D replication in the absence of either G3BP1 or both G3BP1/2. Taken together with our findings with the MEF cells, these results further confirmed an inhibitory function of G3BP1 on MRV replication, and suggest that this inhibition is more impactful on some strains than others.

**Fig. 10.**
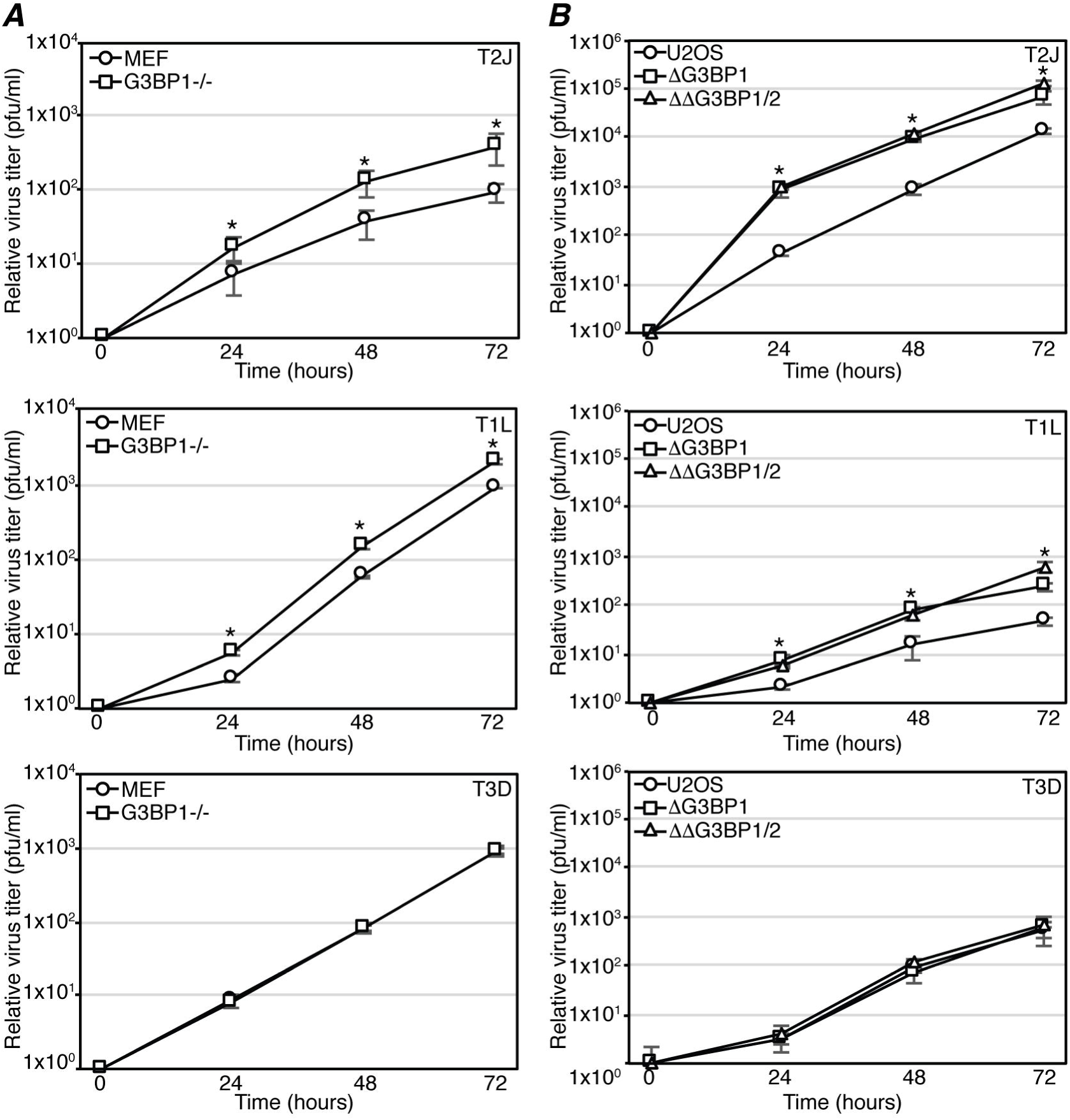
MRV growth is inhibited by G3BP. (A) Wildtype and G3BP1 -/- MEFs or (B) Wildtype, ΔG3BP1 and ΔΔG3BP1/2 U2OS cells were infected with T1L, T2J or T3D for 0, 24, 48 or 72 hours. Cell lysates were collected and lysed by three freeze/thaw cycles. Lysed samples were subjected to standard MRV plaque assay on L929 cells. Plaques were counted, and relative increases in virus titer from time zero were calculated. The means and SD of three biological replicates and two experimental replicates are shown. A statistically significant difference for which *p* < 0.05 is indicated with an asterisk (∗).

## DISCUSSION

We have previously reported that cells respond to MRV by inducing SG formation during early phases of infection in an eIF2α phosphorylation-dependent manner, and as viral infection proceeds, SGs are disrupted from most of the infected cells in a process dependent on synthesis of viral RNA and/or proteins (34). In addition to disrupting SGs over time, MRV prevents SG formation in most cells, even in the presence of phosphorylated eIF2α and SG-inducing drugs such as sodium arsenite. This suggests that MRV acts downstream of these cellular stress signals to interfere with SG formation. Based on these data we hypothesized that MRV encodes a viral component that disrupts or interferes with SG formation, either directly or indirectly by interfering with other SG inducing signals or with SG effecter protein aggregation (35). In this work, we show that during MRV infection, G3BP1, a major SG effector protein, associates with the MRV non-structural protein σNS and this association alters the distribution of additional SG associated proteins Caprin1, USP10, TIA-1, TIAR and eIF3b to the periphery of VFs nucleated by μNS. Our data indicates that co-expression of σNS and μNS interferes with the formation of canonical SG formation and leads to the mislocalization of SG-associated proteins away from their normal SG structure to the periphery of VFLs, suggesting that the G3BP1/σNS/μNS interaction is involved in SG disruption during infection. Although it is likely that this association and re-localization of G3BP1 and other SG-associated proteins plays an important role in MRV-induced SG prevention/disruption, whether it is sufficient to fully interfere with SG formation and function, or whether other factors are involved in this process during MRV infection remains under investigation. The absence of the phenotype in many cells, even following treatment with puromycin or SA [Fig. 1, Fig. 2, Fig. 9, and our previous studies (34)] suggests that it may be an intermediate step in SG disruption by the virus, with cells in which the phenotype is absent, and SG associated proteins are instead diffusely distributed throughout the cell, representing infected cells in which SGs have already been irreversibly disrupted. Intriguing possible mechanisms and outcomes of disruption that we are currently exploring are whether SGs, which are disassembled by heat shock proteins (58), are disassembled upon arrival at VFs by heat shock proteins known to be recruited to VFs (59), and whether VFL relocalization of SG-associated proteins impacts the activity of PKR and other innate immune response factors known to be regulated via SG association (13, 55).

Recent evidence has shown that translation occurs in MRV VFs and that the cellular translational machinery, including ribosomes and translation initiation factors, are localized to VFs (29). σNS was shown to interact with components of the 40S and 43S pre-initiation complex suggesting a role of σNS in recruiting the cellular translational apparatus to the VFs to facilitate viral protein synthesis (29). G3BP1 has also been shown to interact with the 40S ribosomal subunit via its’ RGG domain to mediate SG assembly (10). Our data shows that σNS interacts with G3BP1 at the periphery of VFLs (Fig. 1-4) and individual deletion of either the RNA binding (RRM) and/or the ribosome binding (RGG) regions of G3BP1 substantially decreases but does not abolish G3BP1 localization around VFLs or its association with σNS (Fig. 7 and 8). This suggests that G3BP1 binding to RNA or to the ribosome may play a role in association with σNS and relocalization of G3BP1 around VFLs, but individually, these interactions are not required for G3BP1/VFL recruitment or G3BP1/σNS association. Deletion of both the RRM and RGG domain appears to result in complete loss of G3BP1 localization to VFLs, but although significantly diminished, this mutant maintains measurable interaction with σNS (Fig. 8). This may be interpreted several ways. First, it is possible that the σNS interaction with G3BP1, although important, is not sufficient for recruitment to VFLs, and that some other aspect of G3BP1 activity that is lost in this mutant is contributing to VFL localization. Secondly, it may simply be that the diminishing strength of the interaction is sufficient to prevent recruitment. Finally, it may also be that the PLA detecting σNS interaction with G3BP1 is simply more sensitive than the VFL recruitment assay. As this mutant maintains interaction with σNS, as well as some other G3BP1 functions (10), we believe it is unlikely that this is simply a result of gross protein misfolding. Similar to what we measured with G3BP1 RNA binding, we also found that deletion of the RNA binding residues 2-11 from the N-terminus of σNS does not inhibit G3BP1/σNS association (Fig. 3) suggesting that σNS binding to RNA is also not necessary for σNS/G3BP1 association. While it appears that the σNS/G3BP1 association does not occur solely via either RNA or ribosomes, at this time, we do not know if the interaction occurs through other proteins in a complex, or if it is a direct association. Recent evidence shows that proteins containing a conserved motif (amino acids FGDF) directly bind G3BP1 to inhibit SG formation (54). However, the σNS protein, regardless of strain, does not contain sequence resembling an FGDF motif, therefore, if a direct interaction between σNS and G3BP1 exists, it is through a different mechanism. Finally, although we have shown that G3BP1/σNS interaction is independent of μNS, and G3BP1 is not recruited to VFLs formed by μNS in the absence of σNS, our current (Fig. 9) and previous data (36) indicate that both σNS and a subset of μNS not localized to VFLs are recruited to SGs that are induced by SA treatment. Hence there remains a possibility that a function of μNS independent of VF formation is involved in recruiting G3BP1 and other SG associated proteins to the VF periphery during MRV infection.

Our data indicates that the percentage of infected cells that exhibit G3BP1 localization around VFs is strain-dependent (Fig. 1). However, in cells transfected with plasmids expressing the μNS and σNS proteins of T1L, T2J and T3D, the percentage of cells in which G3BP1 is localized around VFLs is similar between strains (Fig. 4 and data not shown). This suggests that σNS binding to G3BP1, or a complex containing G3BP1, and bringing associated proteins to the periphery of VFs is a conserved phenotype between the σNS and μNS proteins derived from the three strains of virus. Therefore, the discrepancy in the phenotype seen between MRV strains is likely to be upstream of SG formation, most probably at the level of PKR activation and eIF2α phosphorylation. There are previously defined strain-dependent differences that may be involved in the differences we see in G3BP1 localization. T3D infection prevents PKR activation, 2IF2ɑ phosphorylation. There are previously defined strain-dependent differences that may be involved in the differences we see in G3BP1 localization. T3D infection prevents PKR activation, eIF2α phosphorylation, and subsequent host translational shutoff in infected cells, and T2J, and to a lesser extent, T1L infection is unable to inhibit this translational shutoff (1, 60, 61). This phenotype has been mapped to the dsRNA-binding σ3 protein of MRV and is suggested to occur because of differences in σ3 localization and association with the virus membrane penetration protein μ1 between the virus strains (61). We are currently investigating the possible role of σ3, PKR, and eIF2α phosphorylation in modulating the differences seen in G3BP1 localization during infection by the three MRV strains.

Virus replication assays indicate that MRV strains T1L and T2J, but not T3D, grow better in both the G3BP1-/-MEFs and in ΔG3BP1 or ΔΔG3BP1/2 U2OS cells than in their wildtype counterparts (Fig. 10). These results suggest that G3BP1 has an inhibitory effect on T1L and T2J replication but it does not impact T3D replication. This partially mirrors our results examining the localization of G3BP1 to VFs in infected cells, where T2J exhibits the highest number of cells with localization phenotype, followed by T1L and T3D, which exhibit less detectible G3BP1 localization. It is not clear why there are differences between localization (Fig. 1) and replication assays (Fig. 10), however, they may result from differences in assay sensitivity with replication assays being very sensitive relative to immunofluorescence assays that rely on factors such as antibody sensitivity. We have not yet formally ruled out a direct, SG-independent role for G3BP1 in MRV replication, however, strain-specific differences in replication efficiency in the absence of G3BP1 almost exactly mirror differences in the ability of the strains to prevent PKR activation, eIF2α phosphorylation and host translational shutoff (60-63). As stated above, MRV strain T3D σ3 protein binds dsRNA, preventing PKR activation, eIF2α phosphorylation, and host translational shutoff (1, 61). In the absence of eIF2α phosphorylation, SGs will not form in MRV infected cells (34), hence the absence of G3BP1, and the resultant loss of G3BP1-induced SGs, should not, and does not (Fig 10), have an impact on the replication efficiency of the T3D strain. T2J, and to a lesser extent T1L viruses, on the other hand, are unable to prevent PKR activation, eIF2α phosphorylation, and host translational shutoff that results in the formation of G3BP1-induced SGs (1, 61). Hence, when SGs cannot form in the G3BP1 knock-out strains, T1L and T2J viruses should, and do, replicate better than in the wildtype cells. Moreover, as we have shown that all three viruses are able to interfere with normal SG formation (35), this somewhat modest increase of 5-10 fold is not surprising, as it supports the idea that SGs are inhibitory to MRV replication, but the virus has devised a way to overcome this inhibition. Importantly, these findings also suggest that the formation of G3BP1-induced SGs themselves play an inhibitory role in T1L and T2J replication that is likely a critical downstream component of PKR activation, eIF2α phosphorylation, and host translational shutoff induced by these strains.

## ACKNOWLEDGEMENTS

We thank Richard Lloyd for the G3BP1 -/- MEF cell line and Nancy Kedersha and Paul Anderson for the wildtype, ΔG3BP1 and ΔΔG3BP1/2 U2OS cell lines. We additionally thank Kate Carroll for constructing p-G3BP1/EGFP, and Nancy Kedersha for helpful discussion. This work was supported by the Iowa State University College of Veterinary Medicine.

## REFERENCES

1. Smith JA, Schmechel SC, Williams BR, Silverman RH, Schiff LA. 2005. Involvement of the interferon-regulated antiviral proteins PKR and RNase L in reovirus-induced shutoff of cellular translation. J Virol 79:2240-50.

2. Kedersha N, Anderson P. 2002. Stress granules: sites of mRNA triage that regulate mRNA stability and translatability. Biochem Soc Trans 30:963-9.

3. Buchan JR, Parker R. 2009. Eukaryotic Stress Granules: The Ins and Outs of Translation. Mol Cell 36:932-941.

4. Montero H, Trujillo-Alonso V. 2011. Stress Granules in the Viral Replication Cycle. Viruses-Basel 3:2328-2338.

5. Parker F, Maurier F, Delumeau I, Duchesne M, Faucher D, Debussche L, Dugue A, Schweighoffer F, Tocque B. 1996. A ras-GTPase-activating protein SH3-domain-binding protein. Mol and Cell Biol 16:2561-2569.

6. Annibaldi A, Dousse A, Martin S, Tazi J, Widmann C. 2011. Revisiting G3BP1 as a RasGAP binding protein: sensitization of tumor cells to chemotherapy by the RasGAP 317-326 sequence does not involve G3BP1. PLoS One 6:e29024.

7. Tourriere H, Chebli K, Zekri L, Courselaud B, Blanchard JM, Bertrand E, Tazi J. 2003. The RasGAP-associated endoribonuclease G3BP assembles stress granules. J Cell Biol 160:823-831.

8. Takahashi M, Higuchi M, Matsuki H, Yoshita M, Ohsawa T, Oie M, Fujii M. 2013. Stress Granules Inhibit Apoptosis by Reducing Reactive Oxygen Species Production. Mol and Cell Biol 33:815-829.

9. Qiu YQ, Yang CW, Lee YZ, Yang RB, Lee CH, Hsu HY, Chang CC, Lee SJ. 2015. Targeting a ribonucleoprotein complex containing the caprin-1 protein and the c-Myc mRNA suppresses tumor growth in mice: an identification of a novel oncotarget. Oncotarget 6:2148-63.

10. Kedersha N, Panas MD, Achorn CA, Lyons S, Tisdale S, Hickman T, Thomas M, Lieberman J, McInerney GM, Ivanov P, Anderson P. 2016. G3BP-Caprin 1-USP10 complexes mediate stress granule condensation and associate with 40S subunits. J Cell Biol 212:845-860.

11. Reineke LC, Lloyd RE. 2015. The Stress Granule Protein G3BP1 Recruits Protein Kinase R To Promote Multiple Innate Immune Antiviral Responses. J Virol 89:2575-2589.

12. Matsuki H, Takahashi M, Higuchi M, Makokha GN, Oie M, Fujii M. 2013. Both G3BP1 and G3BP2 contribute to stress granule formation. Genes to Cells 18:135-146.

13. Onomoto K, Jogi M, Yoo JS, Narita R, Morimoto S, Takemura A, Sambhara S, Kawaguchi A, Osari S, Nagata K, Matsumiya T, Namiki H, Yoneyama M, Fujita, T. 2012. Critical role of an antiviral stress granule containing RIG-I and PKR in viral detection and innate immunity. PLoS One 7:e43031.

14. Valadão AL, Aguiar RS, de Arruda LB. 2016. Interplay between Inflammation and Cellular Stress Triggered by Flaviviridae Viruses. Front Microbiol 7:1233.

15. Pham AM, Santa Maria FG, Lahiri T, Friedman E, Marié IJ, Levy DE. 2016. PKR Transduces MDA5-Dependent Signals for Type I IFN Induction. PLoS Pathog 12:e1005489.

16. Miller CL. 2011. Stress Granules and Virus Replication. Future Virol 6:1329-1338.

17. White JP, Lloyd RE. 2012. Regulation of stress granules in virus systems. Trends Microbiol 20:175-83.

18. White JP, Cardenas AM, Marissen WE, Lloyd RE. 2007. Inhibition of cytoplasmic mRNA stress granule formation by a viral proteinase. Cell Host & Microbe 2:295-305.

19. McInerney GM, Kedersha NL, Kaufman RJ, Anderson P, Liljestrom P. 2005. Importance of eIF2a phosphorylation assembly in alphavirus translation and stress granule regulation. Mol Biol Cell 16:3753-3763.

20. Montero H, Rojas M, Arias CF, Lopez S. 2008. Rotavirus infection induces the phosphorylation of eIF2α but prevents the formation of stress granules. J Virol 82:1496-1504.

21. Lindquist ME, Lifland AW, Utley TJ, Santangelo PJ, Crowe JE. 2010. Respiratory syncytial virus induces host RNA stress granules to facilitate viral replication. J Virol 84:12274-84.

22. Clements D, Helson E, Gujar SA, Lee PW. 2014. Reovirus in cancer therapy: an evidence-based review. Oncolytic Virother 3:69-82.

23. Thirukkumaran C, Morris DG. 2015. Oncolytic Viral Therapy Using Reovirus. Methods Mol Biol 1317:187-223.

24. Rhim JS, Mayor HD, Jordan LE. 1962. Cytochemical, fluorescent-antibody and electron microscopic studies on growth of reovirus (echo 10) in tissue culture. Virology 17:342-&.

25. Broering TJ, Parker JSL, Joyce PL, Kim JH, Nibert ML. 2002. Mammalian reovirus nonstructural protein μNS forms large inclusions and colocalizes with reovirus microtubule-associated protein μ2 in transfected cells. J Virol 76:8285-8297.

26. Dales S. 1965. Replication of animal viruses as studied by electron microscopy. Am J Med 38:699-715.

27. Wickner RB. 1993. Double-stranded-rna virus-replication and packaging. J Biol Chem 268:3797-3800.

28. Miller CL, Arnold MM, Broering TJ, Hastings CE, Nibert ML. 2010. Localization of mammalian orthoreovirus proteins to cytoplasmic factory-like structures via nonoverlapping regions of μNS. J Virol 84:867-82.

29. Desmet EA, Anguish LJ, Parker JSL. 2014. Virus-Mediated Compartmentalization of the Host Translational Machinery. Mbio 5(5):e01463-14.

30. Becker MM, Peters TR, Dermody TS. 2003. Reovirus σNS and μNS proteins form cytoplasmic inclusion structures in the absence of viral infection. Journal of Virology 77:5948-5963.

31. Broering TJ, Kim J, Miller CL, Piggott CD, Dinoso JB, Nibert ML, Parker JS. 2004. Reovirus nonstructural protein μNS recruits viral core surface proteins and entering core particles to factory-like inclusions. J Virol 78:1882-92.

32. Miller CL, Broering TJ, Parker JSL, Arnold MM, Nibert ML. 2003. Reovirus σNS protein localizes to inclusions through an association requiring the μNS amino terminus. Journal of Virology 77:4566-4576.

33. Miller CL, Arnold MM, Broering TJ, Eichwald C, Kim J, Dinoso JB, Nibert ML. 2007. Virus-derived platforms for visualizing protein associations inside cells. Mol Cell Proteomics 6:1027-38.

34. Qin QS, Hastings C, Miller CL. 2009. Mammalian Orthoreovirus Particles Induce and Are Recruited into Stress Granules at Early Times Postinfection. Journal of Virology 83:11090-11101.

35. Qin QS, Carroll K, Hastings C, Miller CL. 2011. Mammalian Orthoreovirus Escape from Host Translational Shutoff Correlates with Stress Granule Disruption and Is Independent of eIF2α Phosphorylation and PKR. Journal of Virology 85:8798-8810.

36. Carroll K, Hastings C, Miller CL. 2014. Amino acids 78 and 79 of Mammalian Orthoreovirus protein μNS are necessary for stress granule localization, core protein λ2 interaction, and de novo virus replication. Virology 448:133-45.

37. Zekri L, Chebli K, Tourrière H, Nielsen FC, Hansen TV, Rami A, Tazi J. 2005. Control of fetal growth and neonatal survival by the RasGAP-associated endoribonuclease G3BP. Mol Cell Biol 25:8703-16.

38. Becker MM, Goral MI, Hazelton PR, Baer GS, Rodgers SE, Brown EG, Coombs KM, Dermody TS. 2001. Reovirus σNS protein is required for nucleation of viral assembly complexes and formation of viral inclusions. Journal of Virology 75:1459-1475.

39. Mendez, II, Hermann LL, Hazelton PR, Coombs KM. 2000. A comparative analysis of Freon substitutes in the purification of reovirus and calicivirus. Journal of Virological Methods 90:59-67.

40. Parker JS, Broering TJ, Kim J, Higgins DE, Nibert ML. 2002. Reovirus core protein μ2 determines the filamentous morphology of viral inclusion bodies by interacting with and stabilizing microtubules. J Virol 76:4483-96.

41. Broering TJ, Arnold MM, Miller CL, Hurt JA, Joyce PL, Nibert ML. 2005. Carboxyl-proximal regions of reovirus nonstructural protein μNS necessary and sufficient for forming factory-like inclusions. Journal of Virology 79:6194-6206.

42. Eichwald C, Vascotto F, Fabbretti E, Burrone OR. 2002. Rotavirus NSP5: mapping phosphorylation sites and kinase activation and viroplasm localization domains. J Virol 76:3461-70.

43. Schneider CA, Rasband WS, Eliceiri KW. 2012. NIH Image to ImageJ: 25 years of image analysis. Nat Methods 9:671-5.

44. Furlong DB, Nibert ML, Fields BN. 1988. σ1 protein of mammalian reoviruses extends from the surfaces of viral particles. Journal of virology 62:246-256.

45. Smith JA, Schmechel SC, Raghavan A, Abelson M, Reilly C, Katze MG, Kaufman RJ, Bohjanen PR, Schiff LA. 2006. Reovirus induces and benefits from an integrated cellular stress response. J Virol 80:2019-33.

46. Bounedjah O, Desforges B, Wu TD, Pioche-Durieu C, Marco S, Hamon L, Curmi PA, Guerquin-Kern JL, Piétrement O, Pastré D. 2014. Free mRNA in excess upon polysome dissociation is a scaffold for protein multimerization to form stress granules. Nucleic Acids Res 42:8678-91.

47. Kedersha N, Cho MR, Li W, Yacono PW, Chen S, Gilks N, Golan DE, Anderson P. 2000. Dynamic shuttling of TIA-1 accompanies the recruitment of mRNA to mammalian stress granules. J Cell Biol 151:1257-68.

48. Mollet S, Cougot N, Wilczynska A, Dautry F, Kress M, Bertrand E, Weil D. 2008. Translationally repressed mRNA transiently cycles through stress granules during stress. Mol Biol Cell 19:4469-79.

49. Fabbretti E, Afrikanova I, Vascotto F, Burrone OR. 1999. Two non-structural rotavirus proteins, NSP2 and NSP5, form viroplasm-like structures in vivo. J Gen Virol 80 (Pt 2):333-9.

50. Eichwald C, Rodriguez JF, Burrone OR. 2004. Characterization of rotavirus NSP2/NSP5 interactions and the dynamics of viroplasm formation. Journal of General Virology 85:625-634.

51. Yin HS, Lee LH. 1998. Identification and characterization of RNA-binding activities of avian reovirus non-structural protein σNS. J Gen Virol 79 ( Pt 6):1411-3.

52. Gillian AL, Nibert ML. 1998. Amino terminus of reovirus nonstructural protein σNS is important for ssRNA binding and nucleoprotein complex formation. Virology 240:1-11.

53. Irvine K, Stirling R, Hume D, Kennedy D. 2004. Rasputin, more promiscuous than ever: a review of G3BP. International Journal of Developmental Biology 48:1065-1077.

54. Panas MD, Schulte T, Thaa B, Sandalova T, Kedersha N, Achour A, McInerney GM. 2015. Viral and Cellular Proteins Containing FGDF Motifs Bind G3BP to Block Stress Granule Formation. Plos Pathogens 11(2): e1004659.

55. Reineke LC, Kedersha N, Langereis MA, van Kuppeveld FJM, Lloyd RE. 2015. Stress Granules Regulate Double-Stranded RNA-Dependent Protein Kinase Activation through a Complex Containing G3BP1 and Caprin1. Mbio 6(2):e02486-14.

56. Tourrière H, Gallouzi IE, Chebli K, Capony JP, Mouaikel J, van der Geer P, Tazi J. 2001. RasGAP-associated endoribonuclease G3BP: selective RNA degradation and phosphorylation-dependent localization. Mol Cell Biol 21:7747-60.

57. Taniuchi K, Nishimori I, Hollingsworth MA. 2011. The N-Terminal Domain of G3BP Enhances Cell Motility and Invasion by Posttranscriptional Regulation of BART. Molecular Cancer Research 9:856-866.

58. Rikhvanov EG, Romanova NV, Chernoff YO. 2007. Chaperone effects on prion and nonprion aggregates. Prion 1:217-22.

59. Kaufer S, Coffey CM, Parker JS. 2012. The cellular chaperone hsc70 is specifically recruited to reovirus viral factories independently of its chaperone function. J Virol 86:1079-89.

60. Sharpe AH, Fields BN. 1982. Reovirus inhibition of cellular RNA and protein synthesis: role of the S4 gene. Virology 122:381-91.

61. Schmechel S, Chute M, Skinner P, Anderson R, Schiff L. 1997. Preferential translation of reovirus mRNA by a σ3-dependent mechanism. Virology 232:62-73.

62. Huismans H, Joklik WK. 1976. Reovirus-coded polypeptides in infected cells: isolation of two native monomeric polypeptides with affinity for single-stranded and doublestranded RNA, respectively. Virology 70:411-24.

63. Lloyd RM, Shatkin AJ. 1992. Translational stimulation by reovirus polypeptide σ3: substitution for VAI RNA and inhibition of phosphorylation of the alpha subunit of eukaryotic initiation factor 2. J Virol 66:6878-84.

